# Cryptic diversity arises from glacial cycles in Pacific herring, a critical forage fish

**DOI:** 10.1101/2025.05.02.651951

**Authors:** LE Timm, SA Almgren, JA López, JR Glass

## Abstract

Forage fishes are biological drivers throughout the Pacific Ocean, from the Arctic to nearly subtropical latitudes. As a critical trophic link, the health and stability of Pacific herring (*Clupea pallasii*) populations have implications for other marine species, including several targeted by large, productive fisheries. Previous research has indicated marked divergence between Pacific herring in the Bering Sea and the Gulf of Alaska. Seeking to localize this biogeographic break, we generated low coverage whole genome resequencing data for 120 Pacific herring from seven sites across the northern Gulf of Alaska and the eastern Bering Sea and Aleutian Islands. Single nucleotide polymorphisms across the mitogenome (267) and nuclear genome (∼5.6 million) corroborate a biogeographic break in Pacific herring along the Alaska Peninsula and Aleutian Islands, as far west as Unalaska. We identified two distinct populations: one exists along the northern coasts of the Aleutian Islands and in the eastern Bering Sea; the other occupies the southern edge of the Aleutians and the Gulf of Alaska. Two mitochondrial haplogroups co-occurring at collection sites across the Gulf of Alaska suggest secondary contact between two populations, likely representing glacial refugia. Our results underscore the importance of geological events to contextualize the diversification of forage fish species.

## INTRODUCTION

Forage fishes are small pelagic species that function as linchpins of the trophic web, consuming algae and plankton and serving as abundant prey items for commercially important fishes, marine mammals, and seabirds (Surma et al., 2018a). Though taxonomically diverse, forage fishes are unified by their size and “boom-and-bust” population dynamics (Trochta et al., 2020). Globally, forage fishes are economically valuable and provide a variety of ecosystem services (Pikitch et al., 2012; Konar et al., 2019; Lam et al., 2019; Nissar et al., 2023). Several species also hold important roles in Indigenous cultures (Morin et al., 2023), including Pacific herring (Thornton, 2015; Moss, 2016), eulachon (Beveridge et al., 2020), and smelt (Palmer et al., 2018). Forage fishes are especially vulnerable to rising ocean temperatures: in high latitudes, where waters are warming 2x–4x faster than the global average (Chylek et al., 2022; Rantanen et al., 2022), higher sea temperatures are associated with detrimental effects on growth rate, body size, and nutritional value of forage fish species (Gobler et al., 2018; Hollowed et al., 2012; Robards et al., 2002; von Biela et al., 2019). Despite their ecological, economic, and cultural importance, and their vulnerability to abiotic factors associated with climate change, forage fishes are understudied in the Arctic, with a paucity of genetic data in particular (Timm et al., 2023).

Pacific herring (*Clupea pallasii*) is a key forage fish species driving biological processes throughout the North Pacific Ocean, from the Arctic to nearly subtropical latitudes (Hay & McCarter, 1997). Due to Pacific herring’s role as a link between trophic levels, its population health and stability have critical implications for other marine species. The high energy content of Pacific herring makes it a valuable prey species for marine mammals (Surma et al., 2018a,b), seabirds (Bishop et al., 2015), and piscivorous fishes, including several targeted by large, productive fisheries (e.g., Alaska pollock, Pacific salmons, Pacific halibut, etc.) (Brodeur et al., 2014). Annual Pacific herring spawning events deliver an influx of energy to nearshore environments that attract a taxonomically diverse array of predators while also supporting commercial and subsistence fisheries for herring sac roe (Surma et al., 2021). In Alaska, USA, Pacific herring fisheries are managed in geographically designated units (Woodby et al., 2005) and include subsistence, sac roe, and bait harvests. Historically, the Northeast Pacific herring sac roe fishery has held high commercial value in Japanese markets since the 1970s (Lam et al., 2019), and the traditional harvest of sac roe by Alaska Native Tribes continues as an important component of cultural identity and subsistence fishing (Thornton, 2001, 2015). Sac roe harvest corresponds with spring spawning events, while the food and bait fisheries occur during the rest of the year, targeting non-spawning adult herring and possibly targeting adults from multiple spawning populations, the largest of which is in Togiak, Alaska, USA.

Multiple methodologies have been used to characterize the demographic and population structure of Pacific herring throughout Alaska. Herring in the Bering Sea are longer lived, reach notably larger body sizes, and migrate farther than herring in the Gulf of Alaska and Pacific populations further south (Hay et al., 2008). However, comparisons of other physical characteristics including growth and maturation rates, scale age patterns, fatty acid composition, and otolith microchemistry have been unable to reliably describe population structure within oceanic basins (Wespestad & Barton, 1979; Rowell, 1981; Otis et al., 2010).

Previous efforts to characterize the genetic structure of Pacific herring across their geographic range identified marker-dependent patterns of variation. Allozyme variation indicated the presence of two Pacific herring populations: one occupying the eastern Gulf of Alaska (eGOA) and one inhabiting the Bering Sea and Northwest Pacific, with a break along the Alaska Peninsula (Grant & Utter, 1984). Analysis of the mitochondrial cytochrome *b* gene (cytB) confirmed the Northwest Pacific/Bering Sea population and identified some intrusion of those haplotypes into the eastern Pacific, as far as Haida Gwaii off the coast of British Columbia (Liu et al., 2011). Additionally, two other mitochondrial lineages were found to co-occur and dominate throughout eastern Pacific populations (Liu et al., 2011). Results from the mitochondrial control region closely reflected cytB results, and while analysis of ten microsatellite loci nearly recapitulated the allozyme signal, separation was made between the East Pacific, including Togiak in the eastern Bering Sea, and the Northwest Pacific, with some admixture between the two regions (Liu et al., 2012). The observed mitonuclear discordance was hypothesized to be an outcome of secondary contact between lineages that diverged in vicariance during the mid-Pleistocene, 1.3 mya, when glaciation mediated climate cooling would have driven the species into disjunct ranges (Liu et al., 2011; 2012). Microsatellites and mitochondrial markers did not yield evidence of significant genetic structure within the Bering Sea, however these data did suggest a break within Prince William Sound, differentiating an eastern and western population in the Gulf of Alaska (O’Connell et al., 1998; Wildes et al., 2011; Wildes et al., 2018). Within spawning seasons, observations of isolation-by-distance indicated geographical and seasonal spawn-site fidelity, suggesting geographically limited gene flow (Petrou et al. 2021). Prior to the application of next generation sequencing, molecular investigations of Pacific herring predominantly showed regional differences that did not necessarily correspond with spawning stocks. However, with the increased implementation of next generation sequencing, a different demographic picture is emerging.

Whole genome resequencing methods, specifically, are revolutionizing population genetics by addressing information gaps across the Tree of Life (Funk et al., 2012; Hemmer-Hansen et al, 2014). Big Data enhances our ability to inventory adaptive variation (Nielsen et al., 2009), characterize genetic structure with high precision, and identify genomic features such as sex-determining regions (Hansen et al., 2022) and structural variants (Mérot et al., 2020). Genotype likelihoods calculated from low coverage whole genome resequencing (lcWGS) datasets increase capacity for genome scale analysis of large sample sizes (Lou et al., 2021). During the past decade a suite of computational tools have emerged to analyze genotype likelihood datasets and, consequently, lcWGS has gained popularity for investigating genetic population structure in marine species (Clucas et al., 2019; Howe et al., 2024; Timm & Larson et al., 2024; St. John et al., in press), including a variety of forage fish species (Andersson et al., 2024).

Despite the importance of Pacific herring in the North Pacific, our understanding of current and historical genetic population dynamics within this taxon is hampered by data gaps. Here, we used lcWGS to 1) characterize genetic structure of Pacific herring spawning populations from northeastern Pacific sites spanning both previously hypothesized biogeographic breaks; 2) localize the biogeographic break between basins; and 3) describe mitonuclear discordance in the species.

## METHODS

### Sampling

We sampled 120 herring from seven locations (five spawning and two non-spawning sites) spanning the northern Gulf of Alaska (nGOA) and eastern Bering Sea and Aleutian Islands (eBSAI) (Fig. 1; Table 1, S1). Samples were collected from spawning populations in Cordova, Togiak, Port Moller, and Kodiak Island. Kodiak Island samples were obtained from two different locations: Uganik and Kiliuda. Non-spawning populations were sampled in Constantine Bay and Popof Island. Samples were collected by state and federal agency biologists, in full compliance with applicable laws on sampling from natural populations, and were obtained from the Alaska Department of Fish and Game Gene Conservation Lab archive of tissue samples.

**Figure 1.**
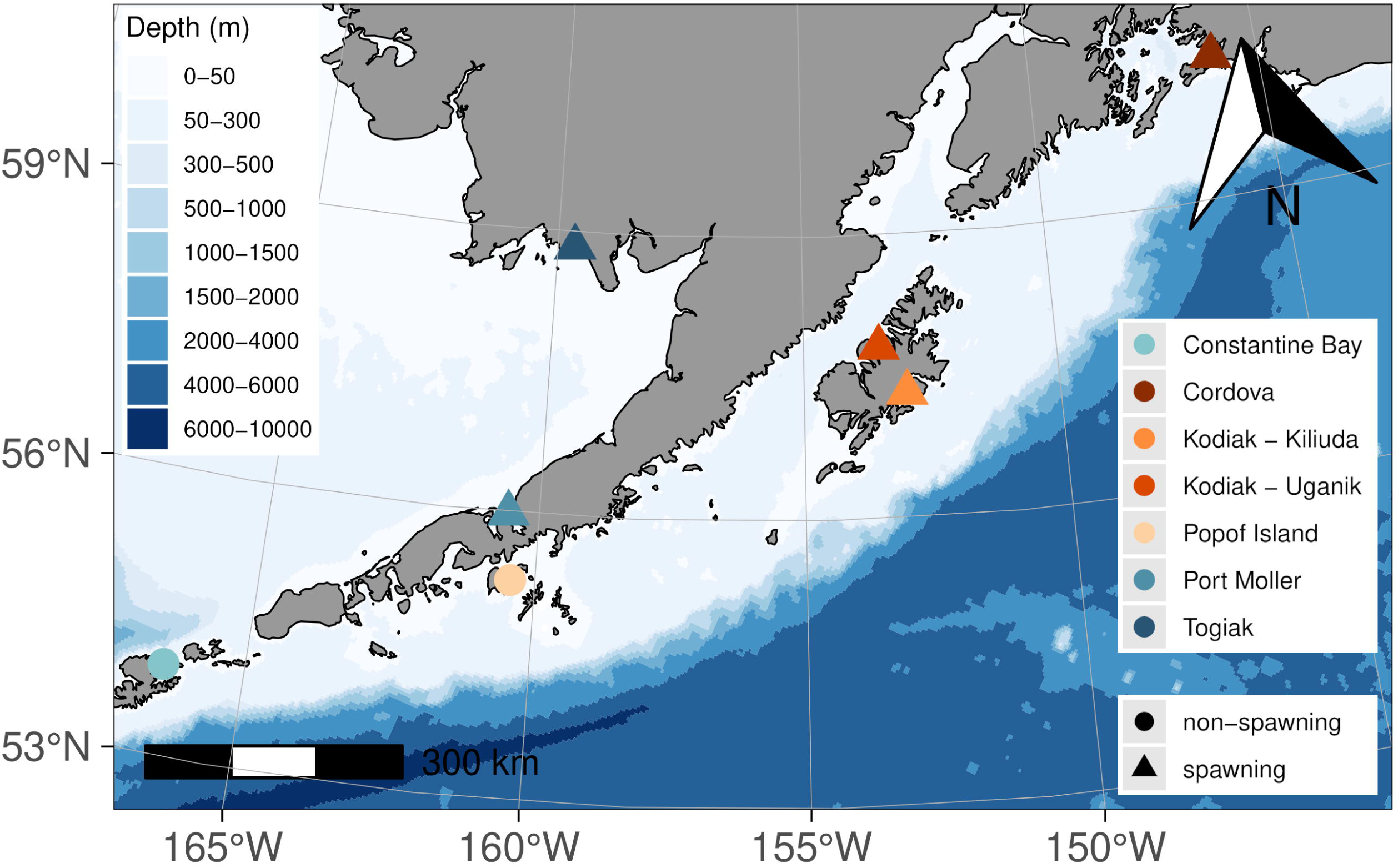
A map of collection locations, colored by geographic region. Spawning status of the collection is indicated by shape.

**Table 1.** Information about the geographic regions included in this study, including the region codes, sample sizes (N), collection date ranges, and diversity values associated with the two-population scenario, including nucleotide diversity (π) and heterozygosity (average individual SFS). Spawning aggregations are in bold.

| Region | N | Collection Date Range | Population | $\pi$ | heterozygosity |
| --- | --- | --- | --- | --- | --- |
| **Cordova** | 20 | March 2023 | nGOA | 0.308 | 0.330 |
| **Kodiak, inside (Uganik)** | 10 | April 2023 |  |  |  |
| **Kodiak, outside (Kiliuda)** | 10 | April 2023 |  |  |  |
| Popof Island | 20 | July 2023 |  |  |  |
| **Port Moller** | 20 | May 1996 | eBSAI | 0.300 | 0.273 |
| Constantine Bay | 20 | July 1999 |  |  |  |
| **Togiak** | 20 | May 2022 |  |  |  |

### Whole genome library preparation and sequencing

Genomic DNA was extracted from tissue samples using Qiagen PureGene Tissue kit following the manufacturer’s protocol (Qiagen, Maryland, United States). Genomic DNA was run on 1% agarose gels to assess quality and quantified with a Qubit 4.0 fluorometer using a dsDNA broad range assay kit (Thermo-Fisher Scientific, United States). Library preparation and sequencing was conducted at the Oklahoma Medical Research Foundation NGS Core (Oklahoma City, OK, United States). Whole genome libraries were constructed using an xGen DNA EZ Library Prep Kit, paired with xGen Normalase UDI primers (Integrated DNA Technologies, Iowa, United States) and KAPA Pure Beads (Roche Molecular Systems, United States) in preparation for paired end sequencing of 150bp. Samples from Togiak and Cordova were sequenced on an Illumina NovaSeq 6000 S4 at up to 6X coverage. Samples from remaining locations were sequenced on an Illumina NovaSeqX with up to 4X coverage.

### Data filtering, alignment, and calculation of genotype likelihoods

Quality of raw reads was ascertained with FastQC v0.12.0 (Andrews, 2010) and aggregated with multiQC v1.19 (Ewels et al., 2016). The TRIMMOMATIC v0.39 (Bolger et al., 2014) *ILLUMINACLIP* tool was used to identify and remove adapters and Illumina-specific sequences (*2:30:10:1:true*), only retaining reads >40bp post-trimming (*MINLEN:40*). We clipped polyG tails using *trim_poly_g* (*-L -A –cut_right*), implemented in fastp v0.23.2 (Chen et al., 2018), to remove a known artefact of two-dye sequencing chemistries.

The Atlantic herring (*Clupea harengus*) reference genome (GCF_900700415.2; Petterson, 2019) was indexed with *index* in BWA v0.7.17 (Li & Durbin, 2009) to facilitate alignment of raw reads to the reference genome. Following trimming and clipping, paired reads were aligned to the indexed reference genome with BWA’s *mem* algorithm. We marked shorter split hits as secondary (*-M*), to ensure alignment results were compatible with downstream filtering programs. Individual alignments were sorted by coordinates with the *sort* tool in samtools v1.19 (Danecek et al., 2021) before removing duplicate reads arising from PCR with the *MarkDuplicates* tool in Picard v2.26.6. Next, overlapping sequence ends were clipped for each mapped read pair with the bamutil v1.0.15 (Table S2) *clipOverlap* tool. Finally, we calculated sequencing depths across the genome of every individual using the *depth* tool in samtools and estimated the mean sequencing depth for each individual using a custom Python script (Table S2). All individuals had a mean sequencing depth >1x and were included in genotype likelihood calculations (Lou et al., 2021).

Genotype likelihoods were estimated for single nucleotide polymorphisms (SNPs) with the samtools model (*-GL 1*) as implemented in ANGSD v0.940 (Korneliussen et al., 2014). For a site to be considered polymorphic, the minimum and maximum depth thresholds were set to the number of individuals in the dataset (*-setMinDepth 120*) and 5x the number of individuals in the dataset (*-setMaxDepth 600*), respectively. Major and minor alleles were inferred from genotype likelihoods (*-doMajorMinor 1*). Candidate SNPs were removed from downstream analysis when: sequencing quality or mapping quality scores were <15 (*-minQ 15 -minMapQ 15*); the *p*-value for polymorphism was greater than 10^-10^ (*-SNP_pval 1e-10*); and the minor allele frequency (MAF) was < 0.05 (*-minMaf 0.05*). Paralogous loci were identified by reading SNP-wise coverage statistics from samtools’ *mpileup* tool into ngsParalog v1.3.4 (Table S2). First, SNPs from reads marked as unmapped or duplicated were disregarded and base and mapping quality thresholds were explicitly set to 0 (*-q 0 -Q 0 –ff UNMAP,DUP*). Mpileup results were piped directly to ngsParalog to calculate the likelihood ratio that each SNP was mismapped to a paralogous site. A Χ^2^ test and Bonferroni correction were used to identify and exclude SNPs with adjusted *p*-values > 0.05.

### Inference of population genetic structure with nuclear data

Genetic variability and structure across the whole dataset was examined with principal components (PCA) and admixture analysis on genome-wide SNPs. We calculated a covariance matrix from SNP genotype likelihoods in PCAngsd v0.99 (Meisner & Albrechtsen, 2018). The minimum average partial test was used to determine the number of eigenvalues to retain (*-e*). We generated eigenvectors by decomposing the covariance matrix with *eigen*, as implemented in R v4.2.0 (R Core Team, 2022). Results from PCA were first used to ascertain whether combining data from two sequencing runs introduced genetic structure artefacts (Lou et al., 2022). After estimating the position of each individual along PC1, we performed a linear regression analysis to estimate correlation between PC1 position and sequencing depth. Based on minimum average partial test results, we tested *K* = 1–7 using three replicates in NGSadmix (Skotte et al., 2013) and the *K* with highest log likelihood was deemed optimal.

### Differentiation, diversity, and the identification of genomic outlier regions

The Atlantic herring reference genome was indexed with *faidx* in samtools to facilitate site frequency spectra estimation in ANGSD. This reference genome was used as the ancestral genome (*-anc* same as *-ref*) in the folded site allele frequency likelihood (SAF) calculation for each region with *doSaf* in ANGSD. Parameters set for genotype likelihood calculation were used for SAF calculation, with depth thresholds tuned to reflect the sample size of each region. Regional SAFs were used to estimate the 2D site frequency spectrum (SFS) and pairwise *F_ST_* with *realSFS* and *realSFS fst*, respectively, in ANGSD. SAFs were also used to calculate diversity statistics (π and heterozygosity) for each population using *realSFS saf2theta* and *thetaStat* in ANGSD.

Statistical significance of pairwise *F_ST_* values was determined with an individual-based permutation test, as implemented in Timm & Larson et al. (2024). Holding population number and sample sizes constant, every permutation randomly assigned individuals to populations, without replacement, and calculated weighted *F_ST_*. Fifty permutations were completed to generate a distribution of weighted *F_ST_* values. Using a custom Python script (Table S2), the mean of this distribution was calculated and used to estimate the cumulative distribution function (CDF) of the *F_ST_* values under an exponential distribution with *p*-value = 1 – CDF (Elhaik, 2012). *P*-values less than 0.05 were deemed statistically significant.

To analyze population differentiation across the genome, pairwise *F_ST_* was estimated between groups and collection regions for every SNP. Site allele frequency likelihoods were used to estimate 2D SFS and calculate *F_ST_* by-site (*realSFS fst stats2*). By-site *F_ST_* values were visualized in Manhattan plots. Spans of elevated *F_ST_* were further investigated with a local score approach (Fariello et al., 2017; Andrews et al., 2023) prior to designation as an outlier region. For each group, SNPs were filtered by MAF (>0.05), mapping quality (>15), and depth (n ≤ depth ≤ n * 5, where n is the number of samples in the group) prior to counting alleles for each SNP in each group *(angsd -do Counts 1 -dumpCounts 3*). Targeting SNPs from both groups, we used Fisher’s Exact Test to identify SNPs with allele frequencies that differed significantly from the background allele frequency distribution for each group-pair. Following Fariello et al. (2017), we analyzed the proximity of SNPs with significant Fisher’s Exact Test results, setting a smoothing parameter ξ= 2. Chromosome-specific significance thresholds were calculated with ɑ = 0.01 and genomic regions exceeding the corresponding significance threshold were designated as outlier regions. To qualify as a genomic region of interest, the loci needed to be 1) under selection (identified by local score analysis) and 2) differentiating between groups (contain SNPs with *F_ST_* > 0.50).

### Inference of population genetic structure with mitogenomic data

As mitochondrial SNPs had sufficient sequencing depth to confidently call genotypes (>10x) and a species-specific mitochondrial genome was available, mitogenomic haplotypes were analyzed separately from nuclear SNPs. Quality filtering and data assembly followed the process described above, though trimmed and clipped reads were aligned to the Pacific herring mitochondrial genome (NC_009578.1; Lavoue et al., 2007). Genotype likelihood data were converted to a fasta file with a custom Python script (Table S2), coding indeterminate genotype likelihoods (<0.99) as missing data. We also removed any sites for which genotype likelihood was >0.99 for the heterozygous state, as that is impossible in the haploid mitogenome. Quality checks were performed and individuals with >10% missing data (n = 13) were removed from the alignment using Geneious Prime v2024.0.5.

Population genomic analysis of mitochondrial SNPs was performed in R. PCA was conducted with *glPCA* from *adegenet* v2.1.5 (Table S2). We conducted an Analysis of Molecular Variance (AMOVA) with *poppr.amova* and calculated pairwise *F_ST_* with *pairwise.WCfst* in *poppr* v2.9.3 (Table S2). Statistical significance was calculated with the individual permutation test implemented in *test.between* in *hierfstat* v0.5-10, specifying 1000 permutations (Table S2).

### Localizing the biogeographic break

We employed an isolation-by-distance (IBD) approach to localize the biogeographic break identified by previous studies, comparing genetic distance (*F_ST_* / (1 - *F_ST_*)) to geographic distance (km) between collection sites, the latter of which was ascertained by plotting the shortest distance between collection sites that did not cross land. We assigned collection sites to groups previously identified by Liu et al. (2011, 2012). Data points were categorized by whether the corresponding pair of collection sites were in physical proximity and thus represented one group or two. For example, Constantine Bay and Port Moller are both located in the eBSAI, so the corresponding pairwise comparison was categorized as “within group,” whereas Togiak and Cordova are located within the eBSAI and nGOA, respectively, so this pairwise comparison was categorized as “between groups.” Given the disparities between nuclear and mitogenomic signals, IBD analysis was conducted on both datasets.

To determine whether the groups described in previous research were also identifiable in whole genome data, we tested for a biogeographic break between the Northeast Pacific and the Bering Sea (Grant & Utter, 1984; Liu et al., 2011) with a Wilcoxon-Mann-Whitney Test comparing *F_ST_* values within groups versus between groups.

### Mitonuclear discordance

Putatively discordant mitochondrial signal was tested for in the nuclear data. We estimated pairwise *F_ST_* from the nuclear SNPs when individuals were categorized according to their mitochondrial haplotype. We also generated a Manhattan plot of *F_ST_* values genome-wide under the same categorization. Finally, we used the local score approach described above to identify proximal SNPs whose allele frequencies significantly differed from the background distribution.

### Computational resources

All plots were generated and several analyses were conducted in R, using packages detailed and cited in Table S2. Figures were prepared for publication in GIMP (GNU Image Manipulation Program). Detailed scripts for data assembly, analysis, and visualization can be found at https://github.com/letimm/pacific-herring_lcWGS.

## RESULTS

### Sequencing and genotype likelihood estimation

Six geographic regions were represented by twenty individuals each, totaling 120 samples (Table 1, S1). Sequencing yielded 1,799,068,017 raw paired end sequences (approximately 14,992,233 sequences per individual). All individual alignments had an estimated mean sequencing depth >1x, with an overall mean of 3.22x (standard deviation = 0.54). Initially, 5,770,843 SNPs were genotyped across all individuals, but 180,993 SNPs (3.1%) were classified as paralogous and removed, resulting in a final set of 5,589,850 SNPs.

Mean sequencing depth estimated for alignment to the *Clupea pallasii* mitogenome was 412.35x (standard deviation = 347.40) across 120 individuals. After removing 10 paralogous SNPs, 267 SNPs were converted from beagle format to FASTA format.

### Inference of population genetic structure with whole genome data

Principal component analysis (PCA; Fig. 2A, Fig. S1A) revealed two primary clusters differentiated along PC1 (24.26% variance) that corroborated the split between the Gulf of Alaska and Bering Sea: one cluster contained individuals from Cordova, Kodiak-Kiliuda, Kodiak-Uganik, and Popof Island (the nGOA population) and the other cluster contained samples from Constantine Bay, Togiak and Port Moller (the eBSAI population). Two individuals from Togiak fell between the eBSAI and nGOA clusters (Fig. 2A). Aside from these individuals, the eBSAI cluster was more tightly grouped; the nGOA cluster exhibited greater spread and, while it contained 10 individuals from Popof Island, half of the individuals from Popof Island spread along PC2 (1.57% variance). Given the high variance explained along PC1, hierarchical PCA was employed to better visualize individuals in PC-space. Individuals in the eBSAI and nGOA clusters were re-analyzed separately with PCA (both including and excluding Popof Island samples). While the eBSAI PCA identified three primary clusters, these clusters did not correspond to collection sites: all three sites were present in every cluster (Fig. 2B). When Popof Island was included in the nGOA PCA, differences between Popof Island and the other sites drove clustering, separating along PC1 (4.28% variance) and PC2 (1.13% variance) (Fig. 2C). When Popof Island was removed, no structure was discernible in the remaining nGOA samples and variance along PC1 and PC2 axes decreased (2.18% and 1.06%, respectively) (Fig. 2D).

**Figure 2.**
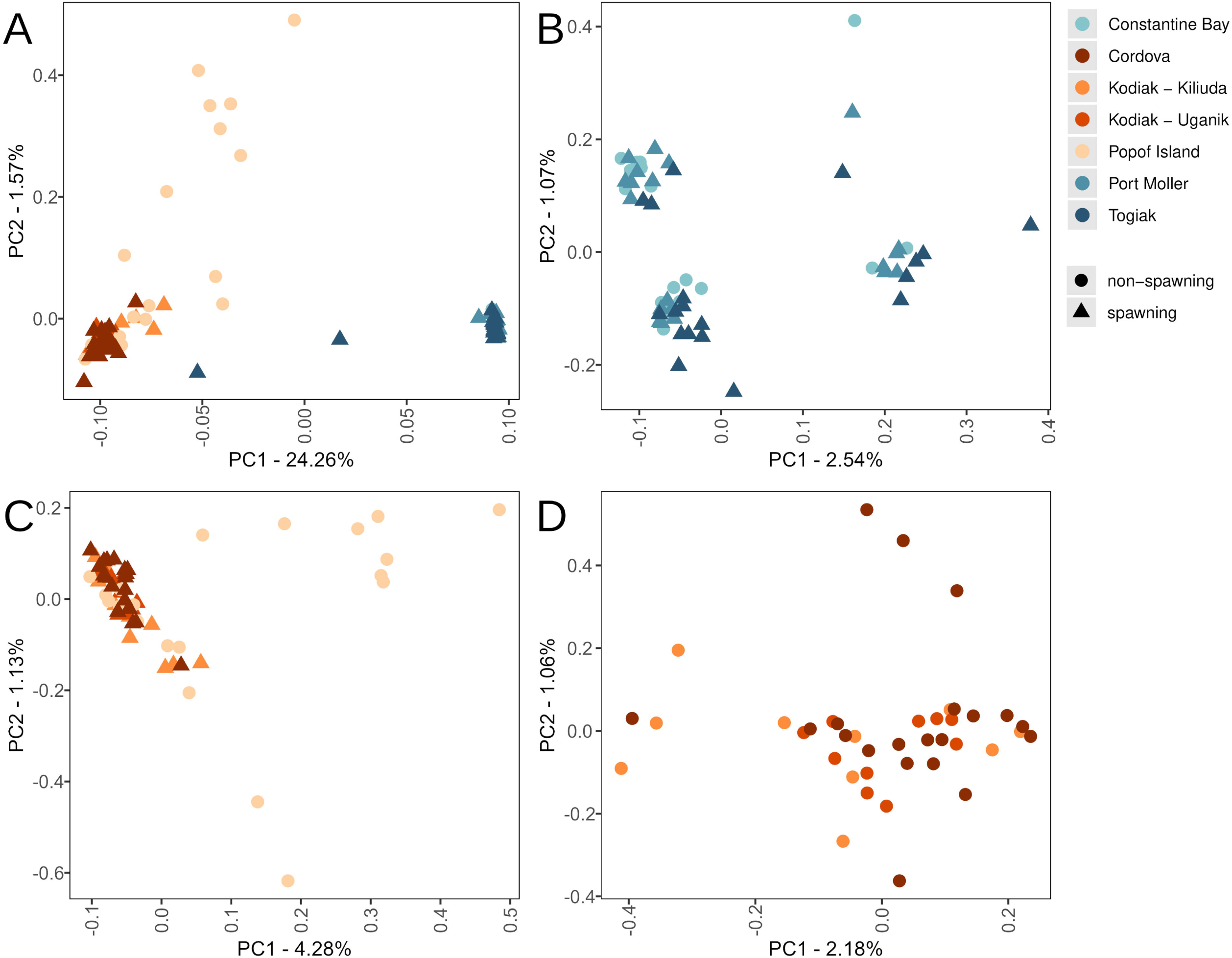
PCAs of A) all samples; B) samples from eBSAI (we denote the upper left cluster as A, the lower left cluster as B, the lower right cluster as C, and the upper right cluster as D); C) samples from nGOA, including Popof Island; and D) samples from nGOA, excluding Popof Island. Color indicates collection region and shape reflects spawning status.

The results of individual admixture proportion analysis largely agreed with the results of hierarchical PCA (Fig. 3). Testing *K* = 2 to *K* = 7 yielded log likelihood values ranging from 484.38 (*K* = 3) to infinite (*K* = 2, optimal). When *K* = 2, the groups nearly completely recapitulate clusters identified with PCA: the eBSAI group contained all individuals from Constantine Bay, Togiak and Port Moller, with no admixture from the nGOA group (with the exception of two Togiak individuals which, in the PCA, appeared midway between the eBSAI cluster and the nGOA cluster). Similarly, the nGOA group contained all individuals from Cordova, Kodiak-Kiliuda, Kodiak-Uganik, and Popof Island with the greatest admixture with the eBSAI group estimated in individuals from Popof Island. As *K* increased, we did not see further separation of discrete groups associated with collection sites. Rather, additional populations contributed proportions of admixture to eBSAI or nGOA. At *K* = 3, for instance, eBSAI individuals showed variable admixture proportions attributable to two populations.

**Figure 3.**
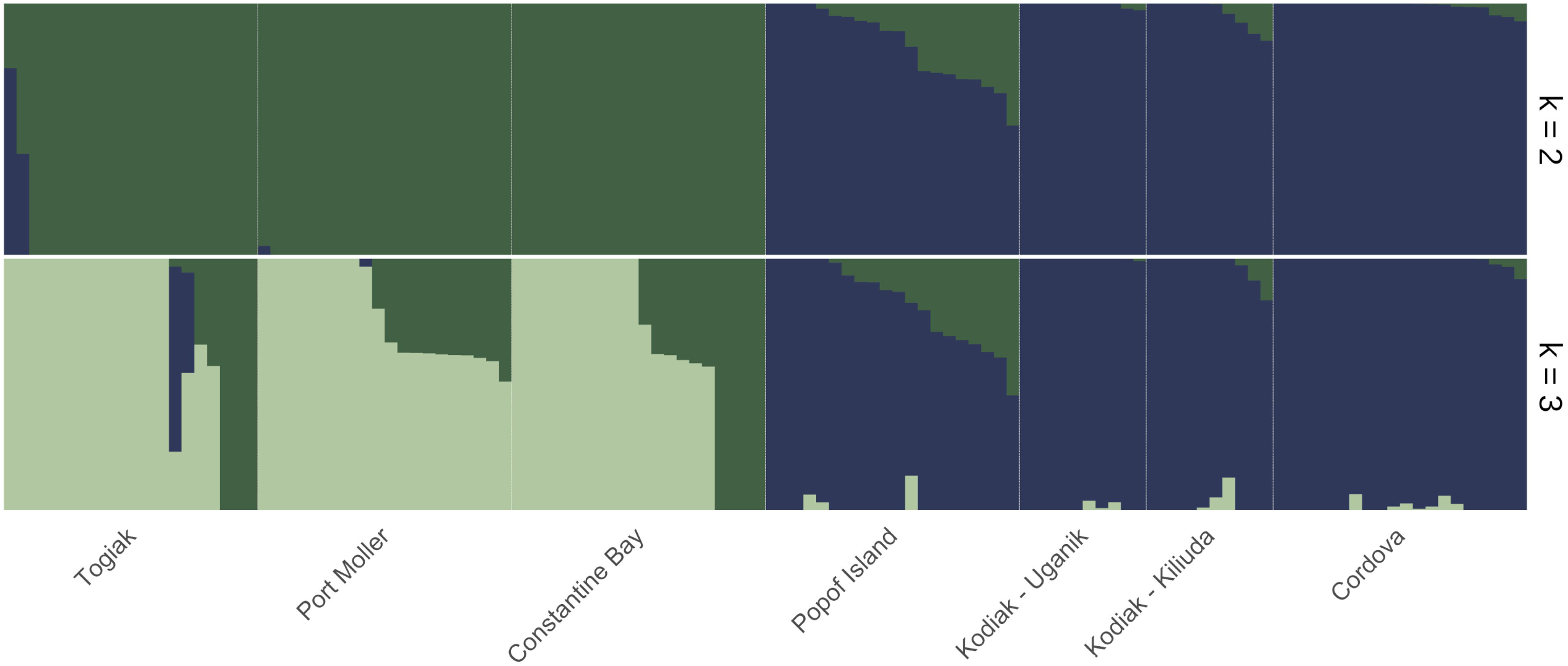
ADMIXTURE results for *K* = 2 and *K* = 3. Log likelihood was highest for *K* = 2, indicating optimality.

### Differentiation, diversity, and the identification of genomic outlier regions

Pairwise *F_ST_* values were calculated between eBSAI and nGOA, with Popof Island included in nGOA (two-population scenario) or defined as a separate population (three-population scenario), as well as between all pairs of collection sites (Table 2). Under the two-population scenario, pairwise *F_ST_* between eBSAI and nGOA was 0.178 (p < 0.001). This value increased when Popof Island was denoted a unique population under the three-population scenario (*F_ST_* = 0.193; p < 0.001; Table S3). Pairwise *F_ST_* was low and insignificant between Popof Island and nGOA (*F_ST_* = 0.022; p > 0.1) and high and significant between Popof Island and eBSAI (*F_ST_* = 0.151; p < 0.001). Because of this, subsequent analyses focused on the two-population scenario: nucleotide diversity (π) was 0.300 in eBSAI and 0.308 in nGOA. Heterozygosity in eBSAI was 0.273 and 0.330 in nGOA.

**Table 2.** Pairwise weighted *F_ST_* values and associated p-values are reported above and below the diagonal, respectively.

|  | Togiak | Port Moller | Constantine Bay | Popof Island | Kodiak - Uganik | Kodiak - Kiliuda | Cordova |
| --- | --- | --- | --- | --- | --- | --- | --- |
| Togiak |  | 0.007 | 0.007 | 0.136 | 0.16 | 0.156 | 0.181 |
| Port Moller | >0.999 |  | 0.006 | 0.128 | 0.146 | 0.144 | 0.169 |
| Constantine Bay | >0.999 | >0.999 |  | 0.126 | 0.142 | 0.14 | 0.167 |
| Popof Island | <0.001 | <0.001 | <0.001 |  | 0.017 | 0.014 | 0.021 |
| Kodiak - Uganik | <0.001 | <0.001 | <0.001 | 0.511 |  | 0.006 | 0.01 |
| Kodiak - Kiliuda | <0.001 | <0.001 | <0.001 | 0.918 | >0.999 |  | 0.011 |
| Cordova | <0.001 | <0.001 | <0.001 | 0.171 | >0.999 | >0.999 |  |

Pairwise *F_ST_* values between collection regions within-population ranged from 0.006 (Kodiak-Uganik versus Kodiak-Kiliuda within nGOA; Constantine Bay versus Port Moller within eBSAI) to 0.021 (Cordova versus Popof Island within nGOA). Intra-population *F_ST_* values were not statistically significant (Table 2). Between populations, pairwise *F_ST_* ranged from 0.126 (Constantine Bay versus Popof Island) to 0.181 (Cordova versus Togiak) and all inter-population comparisons were statistically significant (p < 0.001).

Genome scans of *F_ST_* comparing populations under the two-population and three-population scenarios enabled us to query the data for adaptive loci at unprecedented resolution. However, the differentiation between eBSAI and nGOA was sufficiently high in both scenarios to preclude the identification of adaptive regions (Fig. S2).

Comparisons between Popof Island and the remaining nGOA sites revealed several regions of elevated *F_ST_* (Fig. S3), which were further investigated with the local score approach. Across all pairwise comparisons between collection regions within nGOA, 19 genomic regions in total were identified as potentially under selection: most comparisons only identified 2-3 genomic regions, very rarely on the same chromosome. The Cordova-Popof Island comparison, however, identified seven genomic regions that may be under selection, three of which occur on chromosome 1. Interestingly, the high *F_ST_* block on chromosome 8 was not classified by local score (Fig. S3). In contrast, pairwise comparisons between collection regions in eBSAI only identified five regions with elevated *F_ST_*: two between Constantine Bay and Togiak and three between Port Moller and Togiak, none of which co-occurred on a chromosome (Fig. S4). No genomic regions identified by local score in any of our comparisons included SNPs with *F_ST_* > 0.50.

Within the eBSAI, four clusters, denoted A–D, were identified in the PCA (Fig. 2B), three of which had sufficient sample sizes to perform a genome scan of *F_ST_* and local score analysis (Fig. 4). Local score analysis identified five genomic regions, two between clusters A-C and three between A-B. One genomic region identified by local score also contained SNPs with elevated *F_ST_* (13 SNPs > 0.25; 8 SNPs > 0.50; 1 SNP > 0.75) on chromosome 7 (Fig. 4). When this region was investigated through NCBI’s Genome Viewer, it was found to fall within *tmem8b* (transmembrane protein 8B).

**Figure 4.**
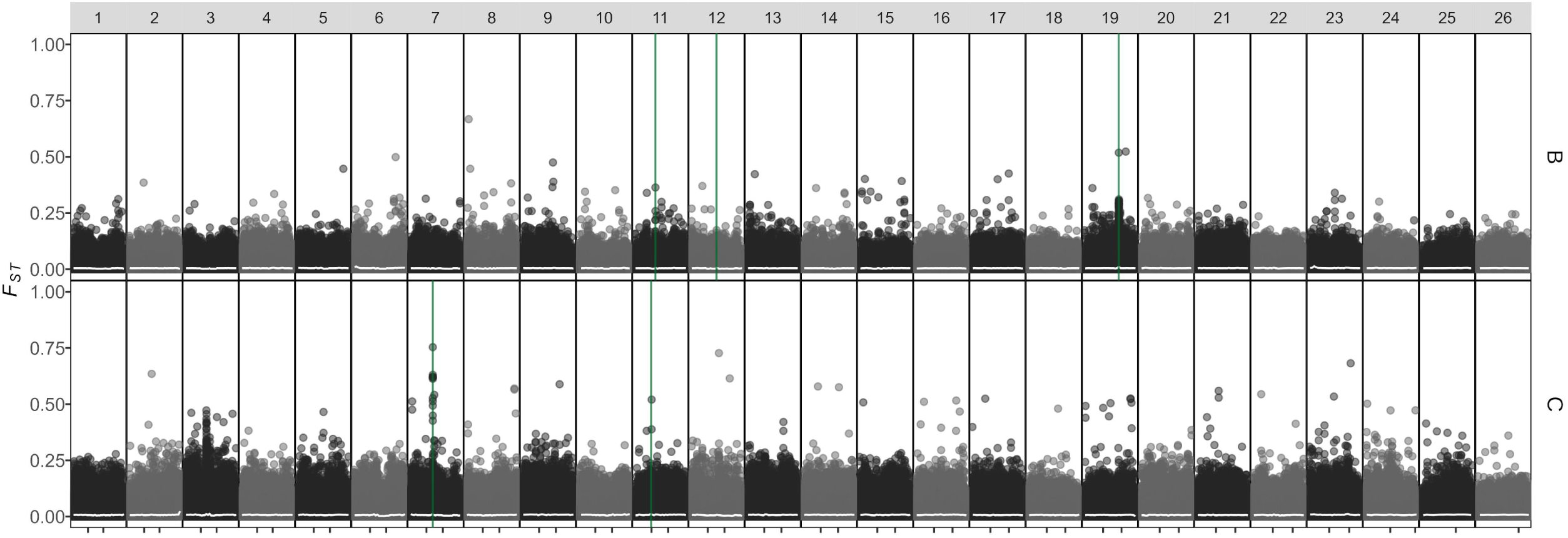
Manhattan plot of *F_ST_* values for all SNPs comparing eBSAI lineage A vs lineage B (top) and lineage C (bottom). Regions identified as under selection by local score analysis are in green. Note the region on chromosome 7, where *F_ST_* and local score identify *tmem8b*.

### Inference of population genetic structure with mitogenomic data

Across the mitogenome, 267 SNPs were genotyped for 107 individuals. The PCA of these data revealed four haplogroups (Fig. 5; Fig. S1B): one was characteristic of the eBSAI (though one of these appears in Kodiak-Uganik and three appear in Popof Island) and three were recovered only in fish from nGOA. The eBSAI haplogroup corresponds to the mitochondrial lineage designated “A” by Liu et al. (2011). The nGOA clusters occupying the upper and lower left hand quadrants are hereafter referred to as GOA1 and GOA2, lineages “B” and “C” in Liu et al. (2011), respectively. Two individuals from Kodiak have mitochondrial genomes belonging to a fourth lineage present in public datasets, but not formally given a designation in prior analysis of mtDNA variation (Liu et al., 2011). A phylogenetic analysis of mitochondrial genomes from representatives of the four distinct haplogroups rooted with *C. harengus* supports a sister group relationship between the eBSAI lineage and a group composed of all other nGOA haplogroups (unpublished data), which are the dominant variants in all examined populations from the Northeast Pacific.

**Figure 5.**
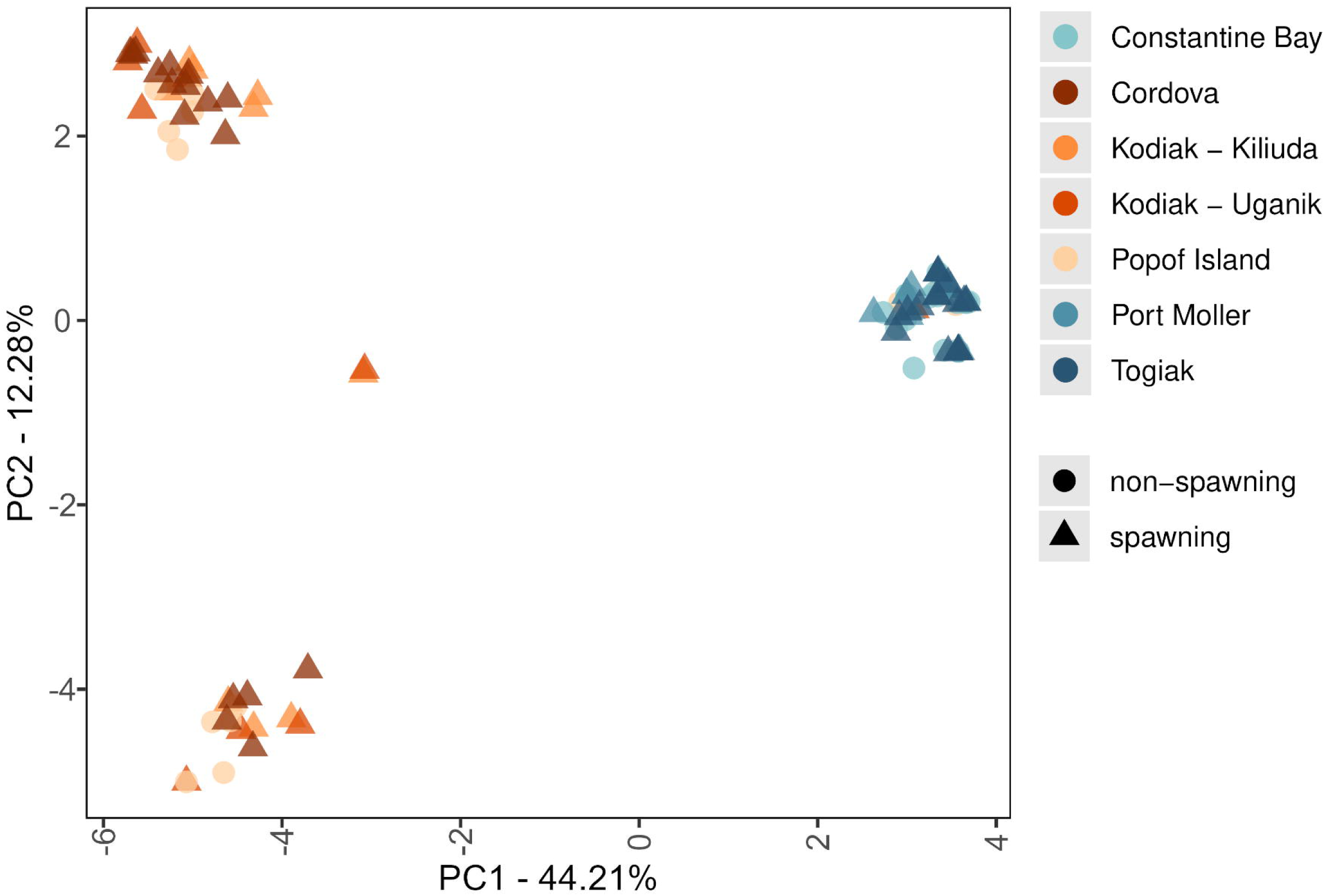
PCA of mitochondrial haplotypes (267 variant sites) from all samples.

The AMOVA found the majority of variance in the data (69.28%) could be attributed to differences between haplogroups (eBSAI versus GOA1 versus GOA2) and the remaining variance was exclusively associated with differences between individuals within collection regions. Differences between collection regions within each haplogroup contributed no variance. All estimates of pairwise *F_ST_* were statistically significant and ranged from 0.697 (eBSAI versus GOA1; eBSAI versus GOA2 = 0.694) to 0.654 (GOA1 vs GOA2). These results agreed with the haplotype analysis: eBSAI had 36 unique haplotypes, seven of which were shared between as many as 11 individuals across three regions (Constantine Bay, Port Moller, and Togiak). GOA1 had 15 unique haplotypes, two of which were shared between two individuals and only one of these spanned collection regions (this haplotype was found in Kodiak-Uganik and Popof Island). GOA2 had 26 unique haplotypes, with only one shared between two individuals in Cordova.

### Localizing the biogeographic break

Analysis of IBD using nuclear SNP data revealed a strong contrast in genetic distances between collection sites corresponding to the “within group” and “between groups” categories. Comparing genetic distance to geographic distance revealed two distinct categories: “within group” genetic distances (comparing two collection regions within nGOA or within eBSAI) were lower than “between group” genetic distances (comparing collection sites between nGOA and eBSAI). Notably, low geographic distance did not correspond to low genetic distance when sites were “between groups”. For example, the geographic distance between Constantine Bay and Port Moller (within group) is nearly equivalent to the geographic distance between Constantine Bay and Popof Island (between group), but the genetic distance of the latter is 24 times greater: 0.006 and 0.144, respectively. When the “within group” data was further divided into “within nGOA” and “within eBSAI,” there were insufficient data points to test for IBD within each region.

Both nuclear and mitochondrial analyses identified a sharp boundary between the eBSAI and nGOA populations within the 400km between Popof Island and Constantine Bay (Fig. S1). No nGOA mitochondrial haplotypes were recovered in the eBSAI and only a few individuals sampled in the nGOA carried eBSAI mitochondrial haplotypes (Fig. 5). The nuclear genome showed a broadly similar pattern, with the exception of two individuals sampled in Togiak that showed strong evidence of admixture between eBSAI and nGOA nuclear genomes (Fig. 2A, 3).

As our data were not normally distributed, we compared distributions of *F_ST_* values between categories with the Wilcoxon-Mann-Whitney Test. The result of this test indicated significantly different distributions (*p*-value = 0.0001), with *F_ST_* values significantly higher in the “between groups” category.

### Mitonuclear discordance

To investigate the degree of concordance between nuclear and mitochondrial data, we performed a genome scan of *F_ST_* values and local score analysis comparing GOA1 and GOA2 in nuclear data (Fig. S5). While a few SNPs had *F_ST_* values > 0.5, none fell within the two regions identified by local score analysis: one on chromosome 10 and another on chromosome 17.

## DISCUSSION

Following the dispersal of Atlantic herring into the North Pacific, facilitated by glacial melting and high sea levels, we identify strong evidence of a subsequent diversification within Pacific herring. Our results provide the most extensive genetic evidence yet for two highly diverged populations of herring in the northeast Pacific. Nuclear and mitochondrial data corroborate a sharp biogeographic break between Bering Sea and northern Pacific populations north and south of the Alaska Peninsula and Aleutian Islands. Additionally, we identify three differentiated lineages of herring in the eBSAI that do not correspond to geographically isolated spawning populations. Finally, we confirm the existence of two distinct mitochondrial haplogroups co-occurring in the northern Gulf of Alaska, likely an outcome of range shifts associated with changes in habitat availability resulting from global climate cycles.

### The Alaska Peninsula and Aleutian Islands as a biogeographic break

Previously, the hypothesized location of a biogeographic break depended on the genetic marker analyzed: allozyme and mitochondrial data inferred a break along the Alaska Peninsula and Aleutian Islands (Grant & Utter, 1984; Liu et al., 2011). Microsatellite variation inferred a break in the North Pacific, differentiating east from west (Liu et al., 2012). Our results concur with the former: we identified a population north of the Aleutian Islands and throughout the eastern Bering Sea and a population south of the Aleutian Islands and across the northern Gulf of Alaska. Our results also indicate that genetic differentiation between the eBSAI and nGOA populations is not limited to a specific genomic region but is genome-wide (Fig. S2). The margins of this division are clearest between Port Moller and Popof Island, two collection sites that are geographically near and exist on the same longitude. The spawning population sampled from Port Moller groups strongly with the other eBSAI collection sites, while Popof Island shows signal that corresponds to the nGOA population. Notably, several individuals from Popof Island cannot confidently be described as belonging to the nGOA or the eBSAI. Samples from Popof Island were not spawning, so these individuals may represent itinerant herring passing through the Aleutian Islands from further south or west.

The break we describe does not appear to be impassable: although we did not test for hybridization directly, we identified two admixed individuals that fell between the eBSAI and nGOA clusters in the PCA (Fig. 2A, 3, S1A). It is notable that these two samples were collected in a spawning aggregation in Togiak and the nearest point of entry from the nGOA, False Pass, is approximately 600km away. These admixed individuals clustered strongly with the eBSAI mitogenomic haplogroup, suggesting intrusion of males from the nGOA into the eBSAI; if females had traversed the break, we would expect these admixed samples to be part of the nGOA haplogroup. Individuals from Constantine Bay (a non-spawning site) exhibited no admixture and only a single sample from Port Moller was admixed with nGOA individuals. Within the nGOA, however, several individuals collected from distinct spawning populations in Kodiak and Cordova indicated admixture with eBSAI individuals.

Pacific herring exhibit a latitudinal cline, spawning earlier in the year at lower latitudes and later in the year at higher latitudes (Haegele & Schweigert, 1985), preventing hybridization through reproductive allochrony. Spawning patterns in Pacific herring (Ljungström et al. 2019) and Atlantic herring (Winters & Wheeler, 1996) are notably dynamic, even amongst geographically proximal populations. Spawn timing of Pacific herring is regulated by annual temperature regimes (Hay 1985) and seasonality (Petrou et al. 2021). Genetic divergence associated with spawn timing has been identified as a driver of population structure in both Atlantic and Pacific herring (Lamichhaney et al., 2017; Petrou et al., 2021). However, herring spawn events have been documented in the northern Gulf of Alaska from February to May, with spawning occurring later in the year as latitude increases (Rounsefell, 1930); herring spawning has been recorded in the Bering Sea from late April to late May (Barton & Wepestad, 1980).

This overlap in spawn timing, in concert with male herring developing earlier than female herring (Hay, 1985), could explain movement of male herring from the nGOA population into eBSAI spawning aggregations and resultant admixture between the populations. The mitogenomic data indicated unidirectional straying in the opposite direction, from the eBSAI to the nGOA: four individuals from the eBSAI haplogroup were collected from two nGOA sites (Fig. S1B). Three of these individuals were collected from Popof Island. This is unsurprising, given the close geographic proximity to Port Moller, an eBSAI spawning population, as well as the non-spawning status of the Popof Island collection. Samples collected from Popof Island may have been passing through the region, rather than returning to spawn. The fourth stray, however, was collected between Kodiak Island and the Alaska Peninsula. This individual had a mitogenomic signal corresponding to the eBSAI, but clustered with nGOA in the nuclear PCA. This was surprising, given that the Alaskan Stream, the predominant oceanographic feature of the region, flows westward (Favorite, 1967), penetrating to the benthos (Warren & Owens, 1985). However, an eastward jet has been described flowing in the deep water south of the Aleutian Islands, north of the Alaskan Stream (Reed, 1969, 1970; Warren & Owens, 1988, Owens & Warren, 2001), which may explain how an individual from the eBSAI haplogroup strayed so far into the nGOA.

Our results indicate this biogeographic break maintains genetic distinctiveness of populations despite some permeability of individuals. Greater unidirectional flow exhibited in mitogenomic data suggests females from the eBSAI may disperse into the nGOA more frequently or with greater success than males and may reflect a general tendency for female herring to move further than males. Pacific herring are hypothesized to exhibit seasonal migrations to spawn and feed in deeper waters of the central Bering Sea, the patterns of which can be highly variable and dependent on environmental conditions such as sea-ice extent and sea surface temperature (Tojo et al., 2007). Dispersal patterns of Pacific herring have been characterized to some degree, but not by sex. However, female herring mature slightly later than males (Hay, 1985) and may use the additional maturation time to migrate further.

### Adaptation and cryptic diversity with gene flow in the eastern Bering Sea

Analyses of genetic differentiation suggest high levels of gene flow between spawning populations within the eBSAI and nGOA. However, we identified at least three sympatric lineages within the eBSAI, suggesting cryptic diversity in this region. Only one genomic region was identified as differentiating between lineages and as being under selection: this gene region corresponded to *tmem8b*, syn. nasopharyngeal carcinoma-associated gene 6 (*ngx6*) is under selection and differentiating eBSAI lineages A and C.

In humans, TMEM8B/NGX6 has been identified as a tumor suppressor (Yang, 2000) and its downregulation is associated with several cancers (Zhang et al., 2003; Ma et al., 2005; Su et al., 2010; Lin et al., 2012). When functional, *ngx6* modulates several cell-signaling pathways, including those associated with physiological processes of metabolism and mitosis (MAPK), intracellular signaling (PI3K/AKT), and proinflammatory signaling (NF-κB) (Li et al., 2012). Two isoforms have been described and both share an epidermal growth factor (EGF)-like extracellular domain. TMEM8B-a is 134 amino acids longer than TMEM8B-b and contains seven transmembrane domains compared to TMEM8B-b’s two domains (Wang et al., 2010). High expression of the TMEM8B-b isoform is hypothesized to improve cell adhesion and reduce the capacity to invade other tissues (Peng et al., 2006).

In domesticated sheep, the only non-human species in which *tmem8b* has been studied to date, *tmem8b* was identified as having been under extreme selection pressure (Kijas et al., 2012). Recently, Cinar et al. (2020) associated *tmem8b* genotype with mature weight at 3-4yrs across several sheep breeds, indicating selection for increased growth rate and larger body size during domestication drove selection at a SNP in *tmem8b*. Interestingly, Pacific herring in the Bering Sea are significantly larger than those in the Gulf of Alaska (Hay et al., 2008), and Bering Sea herring reach larger body sizes at higher latitudes (Schweigert et al., 2002). Hay et al. (2008) suggested this phenotypic difference was the result of local adaptation to resource availability and oceanographic regimes.

Because our results are from a single time point, it is not clear whether this adaptive signal is intensifying or diminishing. Either outcome gives rise to additional questions on the evolution of Pacific herring. If the signal is intensifying, what genetic mechanisms (e.g., inversions) are facilitating adaptation with gene flow? If the signal is diminishing, what is preventing it from homogenizing on the timeline we expect (a few generations)? Answers to either question are critical to recapitulating the evolutionary history of high latitude Pacific herring.

### Secondary contact between glacial refugia

Molecular clocks estimate the colonization of Atlantic herring into the Pacific Ocean occurred during the Pliocene, approximately 1.5–3.5 mya (Grant, 1986; Martinez Barrio et al., 2016, Jamsandekar et al., 2024). Northern GOA herring are thought to have diverged from eBSAI herring during the Pleistocene and to have existed in two refugia in the eastern Pacific during the Last Glacial Maximum (LGM, ∼1 mya) (Liu et al., 2011). The published mutation rate for the mitochondrial genome, 1.59% per million years, roughly agrees with this timing, estimating 266 SNPs separating eBSAI from nGOA: we identified 267. During the LGM, herring in the northern Gulf of Alaska presumably moved south and further diverged in vicariance, which we observe as three nGOA haplogroups. Vicariant divergence was followed by secondary contact in the nGOA, where spawning habitat became available approximately 18,000 years ago (Hughes & Gibbard, 2015). Subsequent mixing homogenized nuclear signal in the nGOA, while mitochondrial signal remained intact due to the mitogenomes’ lack of recombination.

Post-glacial contact has been identified as a driver of current genetic population structure in a number of marine species in the North Pacific, including Pacific cod (Grant et al., 1987; Canino et al., 2010a), walleye pollock (Grant et al., 2010), and Atka mackerel (Canino et al., 2010b). Notably, mitonuclear discordance was also identified in Atka mackerel, another forage fish species in the North Pacific. Canino et al. (2010b) describe genetic homogeneity in Atka mackerel spanning from Japan to the western Gulf of Alaska on the basis of nuclear and mitochondrial markers (microsatellites and the control region). However, mitochondrial diversity was much lower than expected, with only three haplotypes recorded from nearly 120 individuals, suggesting a population bottleneck or founder event associated with climate-driven distribution shifts (Canino et al., 2010b).

The results of our study reify the importance of geological events to understanding species diversification in the present. Taken together, results from Atka mackerel and Pacific herring suggest that range shifts associated with global climate cycles must be considered when interpreting present day genetic population structure and mitonuclear discordance. Efforts to understand the complex interactions between environment and evolution in Pacific herring and other forage fish species may also prove informative for predicting the impact of future climate fluctuations on these key trophic links.

## CONCLUSION

We identify two distinct populations of Pacific herring: one occupies the eastern Bering Sea north of the Aleutian Islands and the other exists in the northern Gulf of Alaska, south of the Aleutian Islands. The lack of gene flow between these two populations of Pacific herring is similar to that described between Pacific and Atlantic herring, suggesting nGOA and eBSAI herring may represent distinct species. We hypothesize nGOA herring separated from eBSAI herring during the Pleistocene, but phylogenetic analysis with a molecular clock is necessary to estimate divergence timing. The lack of eBSAI-nGOA admixture is somewhat surprising given that Atlantic-Pacific hybrid herring are known to exist in stable populations in the Barents Sea (Petterson et al., 2023) and Rossfjordvatn Lake in Norway (Strelkov et al., 2025) but may be attributable to reproductive allochrony. Mitonuclear discordance in the nGOA underscores the importance of contextualizing population genetics in terms of the geological history of the region.

## ACKNOWLEDGEMENTS

The authors thank the Alaska Department of Fish and Game for assistance in sample collection, especially Wei Cheng, Jodi Estrada, Heather Hoyt, Heather Scannel, Jennifer Morella, Elisabeth Fox, M. Birch Foster, Sherri Dressel, Ethan Nichols, and Stacy Vega. Gratitude is also extended to several University of Alaska Fairbanks students for their assistance in sample processing: Isabelle Nicolier, Logan Ito, and Jaden Andrew. We are also appreciative of further sampling and processing help from Bill Carter and Hunter Parini. This research was made possible by grant PCCRC 2022-02 from the Pollock Conservation Cooperative Research Center, as well as grant #23-097G from the Pacific States Marine Fisheries Commission. Special thanks is also given to the anonymous donor of the University of Alaska Fairbanks North Gulf of Alaska Applied Research Award.

## DATA ACCESSIBILITY

Raw sequence reads are deposited in the NCBI SRA (BioProject XXXXX). A detailed description of data assembly, filtering, and analysis is available at https://github.com/letimm/pacific-herring_lcWGS.

## AUTHOR CONTRIBUTIONS

SAA, JAL, and JRG designed the research project. LET performed the research and analyzed the data, with input from the other coauthors. LET wrote the manuscript, with assistance from all coauthors.

## SUPPLEMENTAL MATERIALS

Supplemental Table 1.

Metadata associated with sample collections, including the collection date, coordinates, region, and spawning status of the site. Individual size data, including length (in cm) and weight (in grams) are also reported.

Supplemental Table 2. Computational resources and custom scripts.

Supplemental Table 3. Pairwise weighted *Fst* values between regions under the three-population scenario, where Popof Island (PI), northern Gulf of Alaska (nGOA), and the eastern Bering Sea and Aleutian Islands (eBSAI) are analyzed as separate regions. *P*-values are provided below the diagonal.

Supplemental Figure 1. PCAs by collection region for A) nuclear and B) mitogenomic data. Color indicates collection region and shape reflects spawning status.

Supplemental Figure 2. Manhattan plot of *FST* values for all SNPs comparing eBSAI and nGOA.

Supplemental Figure 3. Manhattan plot of *FST* values for all SNPs comparing Popof Island vs Cordova (top), Kodiak - Uganik (middle), and Kodiak - Kiliuda (bottom). Genomic regions identified as under selection by local score analysis are in green.

Supplemental Figure 4. Manhattan plot of *FST* values for all SNPs comparing Togiak vs Constantine Bay (top) and Port Moller (bottom). Regions identified as under selection by local score analysis are in green.

Supplemental Figure 5. Manhattan plot of *FST* values for all nuclear SNPs comparing GOA1 vs GOA2, as identified by mitochondrial haplotype.

## REFERENCES

Andersson, L., Bekkevold, D., Berg, F., Farrell, E. D., Felkel, S., Ferreira, M. S., Fuentes-Pardo, A. P., Goodall, J., & Pettersson, M. (2024). How fish population genomics can promote sustainable fisheries: a road map. Annual Review of Animal Biosciences, 12(1), 1–20. 10.1146/annurev-animal-021122-102933

Andrews, K. R., Seaborn, T., Egan, J. P., Fagnan, M. W., New, D. D., Chen, Z., Hohenlohe, P.A., Waits, L.P., Caudill, C.C., & Narum, S. R. (2023). Whole genome resequencing identifies local adaptation associated with environmental variation for redband trout. Molecular Ecology, 32(4), 800–818.

Andrews, S. (2010). FastQC: A Quality Control Tool for High Throughput Sequence Data [Online]. Available online at: http://www.bioinformatics.babraham.ac.uk/projects/fastqc/

Aphalo, P. (2023). ggpmisc: Miscellaneous Extensions to ’ggplot2’. R package version 0.5.4-1, <https://CRAN.R-project.org/package=ggpmisc>.

Auguie, B., Antonov, A., & Auguie, M. B. (2017). Package ‘gridExtra’. Miscellaneous functions for “grid” graphics, 9.

Beveridge, R., Moody, M., Murray, G., Darimont, C., & Pauly, B. (2020). The Nuxalk Sputc (Eulachon) project: Strengthening Indigenous management authority through community-driven research. Marine Policy, 119, 103971.

Bishop, M. A., Watson, J. T., Kuletz, K., & Morgan, T. (2015). Pacific herring (*Clupea pallasii*) consumption by marine birds during winter in Prince William Sound, Alaska. Fisheries Oceanography, 24(1), 1–13. 10.1111/fog.12073

Bivand, R. (2022). R packages for analyzing spatial data: A comparative case study with areal data. Geographical Analysis, 54(3), 488–518.

Bougeard, S., & Dray, S. 2018. Supervised multiblock analysis in R with the ade4 package. Journal of Statistical Software, 86(1): 1–17. doi:10.18637/jss.v086.i01

Brodeur, R. D., Buchanan, J. C., & Emmett, R. L. (2014). Pelagic and demersal fish predators on juvenile and adult forage fishes in the Northern California Current: Spatial and temporal variations. California Cooperative Oceanic Fisheries Investigation Report, 55, 96–116.

Canino, M. F., Spies, I. B., Cunningham, K. M., Hauser, L., & Grant, W. S. (2010a). Multiple ice-age refugia in Pacific cod, *Gadus macrocephalus*. Molecular Ecology, 19(19), 4339–4351.

Canino, M. F., Spies, I. B., Lowe, S. A., & Grant, W. S. (2010b). Highly discordant nuclear and mitochondrial DNA diversities in Atka mackerel. Marine and Coastal Fisheries, 2(1), 375–387.

Chen, S., Zhou, Y., Chen, Y., & Gu, J. (2018). fastp: an ultra-fast all-in-one FASTQ preprocessor. Bioinformatics, 34(17), i884–i890.

Chessel, D., Dufour, A., & Thioulouse, J. (2004). The ade4 Package – I: One-Table Methods. R News, 4(1): 5–10. https://cran.r-project.org/doc/Rnews/.

Chylek, P., Folland, C., Klett, J. D., Wang, M., Hengartner, N., Lesins, G., & Dubey, M. K. (2022). Annual mean arctic amplification 1970–2020: observed and simulated by CMIP6 climate models. Geophysical Research Letters, 49(13), e2022GL099371.

Cinar, M. U., Mousel, M. R., Herndon, M. K., Taylor, J. B., & White, S. N. (2020). Association of TMEM8B and SPAG8 with mature weight in sheep. Animals, 10(12), 2391.

Clucas, G. V., Lou, R. N., Therkildsen, N. O., & Kovach, A. I. (2019). Novel signals of adaptive genetic variation in northwestern Atlantic cod revealed by whole-genome sequencing. Evolutionary Applications, 12(10), 1971–1987.

Csárdi, G., Hester, J., Wickham, H., Chang, W., Morgan, M., & Tenenbaum, D. (2021). Remotes: R package installation from remote repositories, including’GitHub’. R package version, 2(2), 86.

Danecek, P., Bonfield, J. K., Liddle, J., Marshall, J., Ohan, V., Pollard, M. O., Whitwham, A., Keane, T., McCarthy, S.A., Davies, R.M., & Li, H. (2021). Twelve years of SAMtools and BCFtools. Gigascience, 10(2), giab008.

Dray, S., & Dufour, A. (2007). The ade4 package: implementing the duality diagram for ecologists. Journal of Statistical Software, 22(4), 1–20. doi:10.18637/jss.v022.i04.

Dray, S., Dufour, A., & Chessel, D. (2007). The ade4 Package – II: Two-Table and K-Table Methods. R News, 7(2), 47–52. https://cran.r-project.org/doc/Rnews/.

Elhaik, E. (2012). Empirical distributions of F ST from large-scale human polymorphism data. PloS ONE, 7(11), e49837.

Fariello, M. I., Boitard, S., Mercier, S., Robelin, D., Faraut, T., Arnould, C., Recoquillay, J., Bouchez, O., Salin, G., Dehais, P., Gourichon, D., Leroux, S., Pitel, F., Leterrier, C., & SanCristobal, M. (2017). Accounting for linkage disequilibrium in genome scans for selection without individual genotypes: the local score approach. Molecular Ecology, 26(14), 3700–3714.

Favorite, F. (1967). The Alaskan Stream. International North Pacific Fisheries Commission Bulletin, 21, 1–20.

Funk, W. C., McKay, J. K., Hohenlohe, P. A., & Allendorf, F. W. (2012). Harnessing genomics for delineating conservation units. Trends in Ecology & Evolution, 27(9), 489–496.

Gobler, C. J., Merlo, L. R., Morrell, B. K., & Griffith, A. W. (2018). Temperature, acidification, and food supply interact to negatively affect the growth and survival of the forage fish, Menidia beryllina (Inland Silverside), and Cyprinodon variegatus (Sheepshead Minnow). Frontiers in Marine Science, 5, 86.

Goudet, J. (2005). Hierfstat, a package for R to compute and test hierarchical *F*-statistics. Molecular Ecology Notes, 5, 184–186

Grant, W. S., Spies, I., & Canino, M. F. (2010). Shifting-balance stock structure in North Pacific walleye pollock (*Gadus chalcogrammus*). ICES Journal of Marine Science, 67(8), 1687–1696.

Grant, W. S., Zhang, C. I., Kobayashi, T., & Ståhl, G. (1987). Lack of genetic stock discretion in Pacific cod (*Gadus macrocephalus*). Canadian Journal of Fisheries and Aquatic Sciences, 44(3), 490–498.

Grant, W. S. (1986). Biochemical genetic divergence between Atlantic, *Clupea harengus*, and Pacific, C. pallasi, herring. Copeia, 714–719.

Grant, W. S., & Utter, F. M. (1984). Biochemical population genetics of Pacific herring (*Clupea pallasi*). Canadian Journal of Fisheries and Aquatic Sciences, 41(6), 856–864.

Haegele, C. W., & Schweigert, J. F. (1985). Distribution and characteristics of herring spawning grounds and description of spawning behavior. Canadian Journal of Fisheries and Aquatic Sciences, 42(S1), s39–s55.

Hansen, C. C. R., Westfall, K. M., & Pálsson, S. (2022). Evaluation of four methods to identify the homozygotic sex chromosome in small populations. BMC Genomics, 23(1), 160.

Hay, D. E. (1985). Reproductive biology of Pacific herring (*Clupea harengus pallasi*). Canadian Journal of Fisheries and Aquatic Sciences, 42(S1), s111–s126.

Hay, D. E., & McCarter, P. B. (1997). Larval distribution, abundance, and stock structure of British Columbia herring. Journal of Fish Biology, 51, 155–175.

Hay, D. E., Rose, K. A., Schweigert, J., & Megrey, B. A. (2008). Geographic variation in North Pacific herring populations: Pan-Pacific comparisons and implications for climate change impacts. Progress in Oceanography, 77(2-3), 233–240.

Hemmer-Hansen, J., Therkildsen, N. O., & Pujolar, J. M. (2014). Population genomics of marine fishes: next-generation prospects and challenges. The Biological Bulletin, 227(2), 117–132.

Hijmans RJ, Phillips S, Leathwick J, & Elith J (2021). dismo: Species Distribution Modeling. R package version 1.3-5, <https://CRAN.R-project.org/package=dismo>.

Hollowed, A. B., Barbeaux, S. J., Cokelet, E. D., Farley, E., Kotwicki, S., Ressler, P. H., Spital, C., & Wilson, C. D. (2012). Effects of climate variations on pelagic ocean habitats and their role in structuring forage fish distributions in the Bering Sea. Deep Sea Research Part II: Topical Studies in Oceanography, 65, 230–250.

Howe, N. S., Hale, M. C., Waters, C. D., Schaal, S. M., Shedd, K. R., & Larson, W. A. (2024). Genomic evidence for domestication selection in three hatchery populations of Chinook salmon, *Oncorhynchus tshawytscha*. Evolutionary Applications, 17(2), e13656.

Hughes, P. D., & Gibbard, P. L. (2015). A stratigraphical basis for the Last Glacial Maximum (LGM). Quaternary International, 383, 174–185.

Jamsandekar, M., Ferreira, M. S., Pettersson, M. E., Farrell, E. D., Davis, B. W., & Andersson, L. (2024). The origin and maintenance of supergenes contributing to ecological adaptation in Atlantic herring. Nature Communications, 15(1), 9136. 10.1038/s41467-024-53079-7

Jombart, T. (2008). adegenet: a R package for the multivariate analysis of genetic markers. Bioinformatics, 24(11), 1403–1405.

Jombart, T., & Ahmed, I. (2011). adegenet 1.3-1: new tools for the analysis of genome-wide SNP data. Bioinformatics, 27(21), 3070–3071.

Kałędkowski, D. (2023). runner: Running Operations for Vectors. R package version 0.4.3, https://CRAN.R-project.org/package=runner.

Kamvar, Z. N., Tabima, J. F., & Grünwald, N. J. (2014). Poppr: an R package for genetic analysis of populations with clonal, partially clonal, and/or sexual reproduction. PeerJ, 2, e281.<doi:10.7717/peerj.281>

Kamvar, Z. N., Brooks, J. C., & Grünwald, N. J. (2015). Novel R tools for analysis of genome-wide population genetic data with emphasis on clonality. Frontiers in Genetics, 6, 208. doi:10.3389/fgene.2015.00208, 10.3389/fgene.2015.00208

Kassambara, A. (2020). ggpubr: ’ggplot2’ Based Publication Ready Plots. R package version 0.4.0, https://CRAN.R-project.org/package=ggpubr.

Kijas, J. W., Lenstra, J. A., Hayes, B., Boitard, S., Porto Neto, L. R., San Cristobal, M., Servin, B., McCulloch, R., Whan, V., Gietzen, K, Paiva, S., Barendse, W., Ciani, E., Raadsma, H., McEwan, J., Dalrymple, B., & other members of the International Sheep Genomics Consortium. (2012). Genome-wide analysis of the world’s sheep breeds reveals high levels of historic mixture and strong recent selection. PLoS Biology, 10(2), e1001258.

Konar, M., Qiu, S., Tougher, B., Vause, J., Tlusty, M., Fitzsimmons, K., Barrows, R., & Cao, L. (2019). Illustrating the hidden economic, social and ecological values of global forage fish resources. Resources, Conservation and Recycling, 151, 104456.

Korneliussen, T. S., Albrechtsen, A., & Nielsen, R. (2014). ANGSD: analysis of next generation sequencing data. BMC Bioinformatics, 15, 1–13.

Laakkonen, H. M., Lajus, D. L., Strelkov, P., & Väinölä, R. (2013). Phylogeography of amphi-boreal fish: tracing the history of the Pacific herring *Clupea pallasii* in North-East European seas. BMC Evolutionary Biology, 13, 1–16.

Laakkonen, H. M., Strelkov, P., Lajus, D. L., & Väinölä, R. (2015). Introgressive hybridization between the Atlantic and Pacific herrings (*Clupea harengus* and *C. pallasii*) in the north of Europe. Marine Biology, 162, 39–54.

Lam, M. E., Pitcher, T. J., Surma, S., Scott, J., Kaiser, M., White, A. S., Pakhomov, E.A., & Ward, L. M. (2019). Value-and ecosystem-based management approach: the Pacific herring fishery conflict. Marine Ecology Progress Series, 617, 341–364.

Lamichhaney, S., Fuentes-Pardo, A. P., Rafati, N., Ryman, N., McCracken, G. R., Bourne, C., Singh, R., Ruzzante, D.E. & Andersson, L. (2017). Parallel adaptive evolution of geographically distant herring populations on both sides of the North Atlantic Ocean. Proceedings of the National Academy of Sciences, 114(17), E3452–E3461.

Lavoue, S., Miya, M., Saitoh, K., Ishiguro, N. B., & Nishida, M. (2007). Phylogenetic relationships among anchovies, sardines, herrings and their relatives (Clupeiformes), inferred from whole mitogenome sequences. Molecular Phylogenetics and Evolution, 43(3), 1096–1105.

Li, H., & Durbin, R. (2009). Fast and accurate short read alignment with Burrows–Wheeler transform. Bioinformatics, 25(14), 1754–1760.

Li, Y., Luo, Y., Wang, X., Shen, S., Yu, H., Yang, J., & Su, Z. (2012). Tumor suppressor gene NGX6 induces changes in protein expression profiles in colon cancer HT-29 cells. Acta Biochimica et Biophysica Sinica, 44(7), 584–590.

Lin, Z. F., Shen, X. Y., Lu, F. Z., Ruan, Z., Huang, H. L., & Zhen, J. (2012). Reveals new lung adenocarcinoma cancer genes based on gene expression. European Review for Medical & Pharmacological Sciences, 16(9), 1249–1256.

Liu, J. X., Tatarenkov, A., Beacham, T. D., Gorbachev, V., Wildes, S., & Avise, J. C. (2011). Effects of Pleistocene climatic fluctuations on the phylogeographic and demographic histories of Pacific herring (*Clupea pallasii*). Molecular Ecology, 20(18), 3879–3893.

Liu, M., Lin, L., Gao, T., Yanagimoto, T., Sakurai, Y., & Grant, W. S. (2012). What maintains the central North Pacific genetic discontinuity in Pacific herring? PLoS One, 7(12), e50340.

Ljungström, G., Francis, T. B., Mangel, M., & Jørgensen, C. (2019). Parent-offspring conflict over reproductive timing: ecological dynamics far away and at other times may explain spawning variability in Pacific herring. ICES Journal of Marine Science, 76(2), 559–572.

Lou, R. N., Jacobs, A., Wilder, A. P., & Therkildsen, N. O. (2021). A beginner’s guide to low-coverage whole genome sequencing for population genomics. Molecular Ecology, 30(23), 5966–5993.

Lou, R. N., & Therkildsen, N. O. (2022). Batch effects in population genomic studies with low-coverage whole genome sequencing data: Causes, detection and mitigation. Molecular Ecology Resources, 22(5), 1678–1692.

Ma, J., Zhou, J., Fan, S., Wang, L., Li, X., Yan, Q., Zhou, M., Liu, H., Zhang, Q., Zhou, H., Gan, K., Li, Z., Peng, C., Xiong, W., Tan, C., Shen, S., Yang, J., Li, J., & Li, G. (2005). Role of a novel EGF-like domain-containing gene NGX6 in cell adhesion modulation in nasopharyngeal carcinoma cells. Carcinogenesis, 26(2), 281–291.

Martinez Barrio, A., Lamichhaney, S., Fan, G., Rafati, N., Pettersson, M., Zhang, H. E., Dainat, J., Ekman, D., Höppner, M., Jern, P., Martin, M., Nystedt, B., Liu, X., Chen, W., Liang, X., Shi, C., Fu, Y., Ma, K., Zhan, X., Feng, C., Gustafson, U., Rubin, C.-J., Almén, M. S., Blass, M., Casini, M., Folkvord, A., Laikre, L., Ryman, N., Lee, S. M.-Y., Xu, X., & Andersson, L. (2016). The genetic basis for ecological adaptation of the Atlantic herring revealed by genome sequencing. elife, 5, e12081.

Meisner, J., & Albrechtsen, A. (2018). Inferring population structure and admixture proportions in low-depth NGS data. Genetics, 210(2), 719–731.

Mérot, C., Oomen, R. A., Tigano, A., & Wellenreuther, M. (2020). A roadmap for understanding the evolutionary significance of structural genomic variation. Trends in Ecology & Evolution, 35(7), 561–572.

Morin, J., Evans, A. B., & Efford, M. (2023). The rise of Vancouver and the collapse of forage fish: a story of urbanization and the destruction of an aquatic ecosystem on the Salish Sea (1885–1920 CE). Human Ecology, 51(2), 303–322.

Moss, M. L. (2016). The nutritional value of Pacific herring: An ancient cultural keystone species on the Northwest Coast of North America. Journal of Archaeological Science: Reports, 5, 649–655.

Neuwirth, E. (2022). RColorBrewer: ColorBrewer Palettes. R package version 1.1-3, https://CRAN.R-project.org/package=RColorBrewer.

Nielsen, E. E., Hemmer-Hansen, J., Larsen, P. F., & Bekkevold, D. (2009). Population genomics of marine fishes: identifying adaptive variation in space and time. Molecular Ecology, 18(15), 3128–3150.

Nissar, S., Bakhtiyar, Y., Arafat, M. Y., Andrabi, S., Bhat, A. A., & Yousuf, T. (2023). A review of the ecosystem services provided by the marine forage fish. Hydrobiologia, 850(12), 2871–2902.

O’Connell, M., Dillon, M. C., Wright, J. M., Bentzen, P., Merkouris, S., & Seeb, J. (1998). Genetic structuring among Alaskan Pacific herring populations identified using microsatellite variation. Journal of Fish Biology, 53(1), 150–163. 10.1111/j.1095-8649.1998.tb00117.x

Otis, E. O., Heintz, R., & Maselko, J. (2010). Investigation of Pacific Herring (*Clupea pallasii*) Stock Structure in Alaska Using Otolith Microchemistry and Heart Tissue Fatty Acid Composition. Exxon Valdez Oil Spill Restoration Project Final Report, Restoration Project 070769. Alaska Department of Fish and Game-Division of Commercial Fisheries, National Marine Fisheries Service-Auke Bay Laboratory.

Owens, W. B., & Warren, B. A. (2001). Deep circulation in the northwest corner of the Pacific Ocean. Deep Sea Research Part I: Oceanographic Research Papers, 48(4), 959–993.

Palmer, E., Tushingham, S., & Kemp, B. M. (2018). Human use of small forage fish: Improved ancient DNA species identification techniques reveal long term record of sustainable mass harvesting of smelt fishery in the northeast Pacific Rim. Journal of Archaeological Science, 99, 143–152.

Papastamoulis, P. (2015). label. switching: An R package for dealing with the label switching problem in MCMC outputs. arXiv preprint arXiv:1503.02271.

Paradis, E. (2010). pegas: an R package for population genetics with an integrated–modular approach. Bioinformatics, 26, 419–420. doi:10.1093/bioinformatics/btp696.

Paradis, E. & Schliep, K. (2019). ape 5.0: an environment for modern phylogenetics and evolutionary analyses in R. Bioinformatics, 35, 526–528.

Peng, S. P., Li, X. L., Wang, L., Ou-Yang, J., Ma, J., Wang, L. L., Liu, H. Y., Zhou, M., Tang, Y. L., Li, W. S., Luo, X. M., Cao, L., Tang, K., Shen, S. R., & Li, G. Y. (2007). The role of NGX6 and its deletion mutants in the proliferation, adhesion and migration of nasopharyngeal carcinoma 5-8F cells. Oncology, 71(3-4), 273–281.

Petrou, E. L., Fuentes-Pardo, A. P., Rogers, L. A., Orobko, M., Tarpey, C., Jiménez-Hidalgo, I., Moss, M. L., Yang, D., Pitcher, T. J., Sandell, T., Lowry, D., Ruzzante, D. E., & Hauser, L. (2021). Functional genetic diversity in an exploited marine species and its relevance to fisheries management. Proceedings of the Royal Society B, 288(1945), 20202398.

Pettersson, M. E., Rochus, C. M., Han, F., Chen, J., Hill, J., Wallerman, O., Fan, G., Hong, X., Xu, Q., Zhang, H. and Liu, S., Liu, X., Haggerty, L., Hunt, T., Martin, F. J., Flicek, P., Bunikis, I., Folkvord, A., & Andersson, L. (2019). A chromosome-level assembly of the Atlantic herring genome—detection of a supergene and other signals of selection. Genome Research, 29(11), 1919–1928.

Pettersson, M. E., Fuentes-Pardo, A. P., Rochus, C. M., Enbody, E. D., Bi, H., Väinölä, R., & Andersson, L. (2023). A long-standing hybrid population between Pacific and Atlantic herring in a subarctic fjord of Norway. Genome Biology and Evolution, 15(5), evad069.

Pikitch, E. K., Rountos, K. J., Essington, T. E., Santora, C., Pauly, D., Watson, R., Sumaila, U. R., Boersma, P. D., Boyd, I. L., Conover, D. O., Cury, P., Heppell, S. S., Houde, E. D., Mangel, M., Plagányi, É., Sainsbury, K., Steneck, R. S., Geers, T. M., Gownaris, N., & Munch, S. B. (2014). The global contribution of forage fish to marine fisheries and ecosystems. Fish and Fisheries, 15(1), 43–64.

R Core Team (2021). R: A language and environment for statistical computing. R Foundation for Statistical Computing, Vienna, Austria. https://www.R-project.org/.

Rantanen, M., Karpechko, A. Y., Lipponen, A., Nordling, K., Hyvärinen, O., Ruosteenoja, K., Vihma, T., & Laaksonen, A. (2022). The Arctic has warmed nearly four times faster than the globe since 1979. Communications Earth & Environment, 3(1), 168.

Reed, R. K. (1969). Deep water properties and flow in the central North Pacific. Journal of Marine Research, 27, 24–31.

Reed, R. K. (1970). On the anomalous deep water south of the Aleutian Islands. Journal of Marine Research, 28, 371–372.

Robards, M. D., Rose, G. A., & Piatt, J. F. (2002). Growth and abundance of Pacific sand lance, *Ammodytes hexapterus*, under differing oceanographic regimes. Environmental Biology of Fishes, 64, 429–441.

Rounsefell, G. A. (1930). Biology of the Pacific herring: Spawning. *Contribution to the biology of the Pacific herring*, Clupea pallasii, *and the condition of the fishery in Alaska* (No. 1080; pp. 272-292). U.S. Government Printing Office.

Rowell, K. A. (1981). Feasibility of scale pattern analysis techniques for stock identification and growth of Pacific herring (*Clupea harengus pallasi*) from four spawning locations of the eastern Bering Sea. Proceedings of the Fourth Pacific Coast Herring Workshop.

St. John, C., Timm, L. E., Gruenthal, K. M., & Larson, W. A. (2025). Whole genome sequencing reveals substantial genetic structure and evidence of local adaptation in Alaskan red king crab. Evolutionary Applications, in press.

Schweigert, J., Funk, F., Oda, K., & Moore, T. (2002). Herring size-at-age variation in the North Pacific. PICES Scientific Reports, 20, 47–56.

Skotte, L., Korneliussen, T. S., & Albrechtsen, A. (2013). Estimating individual admixture proportions from next generation sequencing data. Genetics, 195(3), 693–702.

Strelkov, P., Lajus, D., Laakkonen, H., Moum, T., & Väinölä, R. (2025). Identity and relationships of the landlocked Rossfjordvatn hybrid herring, and other intergrades of *Clupea pallasii* and *C. harengus* in Northern Europe. Marine Biology, 172(2), 29.

Su, Z., Wang, X., Shen, S., Wang, L., Li, Y., Li, N., & Li, Z. (2010). Expression of 2 transcripts of NGX6 gene in colorectal cancer and the correlation with carcinoembryonic antigen. Journal of Central South University. Medical Sciences, 35(5), 401–408.

Surma, S., Pitcher, T. J., Kumar, R., Varkey, D., Pakhomov, E. A., & Lam, M. E. (2018a). Herring supports Northeast Pacific predators and fisheries: Insights from ecosystem modelling and management strategy evaluation. PLoS One, 13(7), e0196307.

Surma, S., Pakhomov, E. A., & Pitcher, T. J. (2018b). Energy-based ecosystem modelling illuminates the ecological role of Northeast Pacific herring. Marine Ecology Progress Series, 588, 147–161.

Surma, S., Pitcher, T. J., & Pakhomov, E. A. (2021). Trade-offs and uncertainties in Northeast Pacific herring fisheries: ecosystem modelling and management strategy evaluation. ICES Journal of Marine Science, 78(6), 2280–2297.

Thioulouse, J., Dray, S., Dufour, A., Siberchicot, A., Jombart, T., & Pavoine, S. (2018). Multivariate Analysis of Ecological Data with ade4. Springer. doi:10.1007/978-1-4939-8850-1

Thornton, T. F. (2001). Subsistence in northern communities: Lessons from Alaska. Northern Review, (23).

Thornton, T. F. (2015). The ideology and practice of Pacific herring cultivation among the Tlingit and Haida. Human Ecology, 43, 213–223.

Timm, L. E., Larson, W. A., Jasonowicz, A. J., & Nichols, K. M. (2024). Whole genome resequencing of sablefish at the northern end of their range reveals genetic panmixia and large putative inversions. ICES Journal of Marine Science, fsae070.

Timm, L. E., Tucker, N., Rix, A., LaBua, S., López, J. A., Boswell, K. M., & Glass, J. R. (2023). The untapped potential of seascape genomics in the North Pacific. Frontiers in Conservation Science, 4, 1249551.

Tojo, N., Kruse, G. H., & Funk, F. C. (2007). Migration dynamics of Pacific herring (*Clupea pallasii*) and response to spring environmental variability in the southeastern Bering Sea. Deep Sea Research Part II: Topical Studies in Oceanography, 54(23-26), 2832–2848.

Trochta, J. T., Branch, T. A., Shelton, A. O., & Hay, D. E. (2020). The highs and lows of herring: A meta-analysis of patterns and factors in herring collapse and recovery. Fish and Fisheries, 21(3), 639–662. 10.1111/faf.12452

Vihtakari, M. (2023). ggOceanMaps: Plot Data on Oceanographic Maps using ’ggplot2’. R package version 2.1.12, https://mikkovihtakari.github.io/ggOceanMaps.

von Biela, V. R., Arimitsu, M. L., Piatt, J. F., Heflin, B., Schoen, S. K., Trowbridge, J. L., & Clawson, C. M. (2019). Extreme reduction in nutritional value of a key forage fish during the Pacific marine heatwave of 2014-2016. Marine Ecology Progress Series, 613, 171–182.

Wang, L., Xiang, B., Yi, M., Zhang, W. L., Yang, J. B., Peng, S. P., Li, X. L., & Li, G. Y., (2010). Identification of a new seven-span transmembrane protein: NGX6a is downregulated in nasopharyngeal carcinoma and is associated with tumor metastasis. Journal of Histochemistry & Cytochemistry, 58(1), 41–51.

Warnes, G. R., Bolker, B., & Lumley, T. (2021). gtools: Various R Programming Tools. R package version 3.9.2, https://CRAN.R-project.org/package=gtools.

Warren, B. A., & Owens, W. B. (1985). Some preliminary results concerning deep northern-boundary currents in the North Pacific. Progress in Oceanography, 14, 537–551.

Warren, B. A., & Owens, W. B. (1988). Deep currents in the central subarctic Pacific Ocean. Journal of Physical Oceanography, 18(4), 529–551.

Wespestad, V. G., & Barton, L. H. (1979). Distribution and migration and status of Pacific herring. Northwest and Alaska Fisheries Center, National Marine Fisheries Service, National Oceanic and Atmospheric Administration.

Wickham, H. (2007). Reshaping Data with the reshape Package. Journal of Statistical Software, 21(12): 1–20. http://www.jstatsoft.org/v21/i12/.

Wickham, H. (2016). ggplot2: Elegant Graphics for Data Analysis. Springer-Verlag New York. ISBN 978-3-319-24277-4, https://ggplot2.tidyverse.org.

Wickham, H. (2022). stringr: Simple, Consistent Wrappers for Common String Operations. R package version 1.5.0, https://CRAN.R-project.org/package=stringr.

Wickham, H., Averick, M., Bryan, J., Chang, W., McGowan, L. D., François, R., Grolemund, G., Hayes, A., Henry, L., Hester, J., Kuhn, M., Pedersen, T. L., Miller, E., Bache, S. M., Müller, K., Ooms, J., Robinson, D., Seidel, D. P., Spinu, V., Takahashi, K., Vaughan, D., Wilke, C., Woo, K., & Yutani, H. (2019). Welcome to the tidyverse. Journal of Open Source Software, 4(43): 1686. 10.21105/joss.01686.

Wickham, H., François, R., Henry, L., Müller, K., & Vaughan, D. (2023). dplyr: A Grammar of Data Manipulation. R package version 1.1.4, https://github.com/tidyverse/dplyr, https://dplyr.tidyverse.org.

Wickham, H., Bryan, J. (2023). readxl: Read Excel Files. R package version 1.4.3, https://CRAN.R-project.org/package=readxl.

Wickham, H., & Seidel, D. (2023). Scales: Scale functions for visualization. 2020. R package version, 1(1).

Wildes, S. L., Vollenweider, J. J., Nguyen, H. T., & Guyon, J. R. (2011). Genetic variation between outer-coastal and fjord populations of Pacific herring (*Clupea pallasii*) in the eastern Gulf of Alaska. Fishery Bulletin, 109(4).

Wildes, S. L., Nguyen, H., & Guyon, J. R. (2018). Genetic stock structure of herring in Prince William Sound. Exxon Valdez Oil Spill Trustee Council.

Winters, G. H., & Wheeler, J. P. (1996). Environmental and phenotypic factors affecting the reproductive cycle of Atlantic herring. ICES Journal of Marine Science, 53(1), 73–88.

Woodby, D., Carlile, D., Siddeek, S., Funk, F., & Clark, J. H. (2005). Commercial fisheries of Alaska. Alaska Department of Fish and Game, Division of Commercial Fisheries. Special Publication No. 05-09.

Yang, J. (2000). Refined localization and cloning of a novel putative tumor suppressor gene associated with nasopharyngeal carcinoma on chromosome 9p21-22. Chinese Journal of Cancer, 19(1), 6–9.

Zhang, X. M., Sheng, S. R., Wang, X. Y., Wang, J. R., & Li, J. (2003). Expression of tumor related genes NGX6, NAG-7, BRD7 in gastric and colorectal cancer. World Journal of Gastroenterology: WJG, 9(8), 1729.

