## Supplementary figures and images for "Cryptic diversity arises from glacial cycles in Pacific herring, a critical forage fish"

### Fig. S1

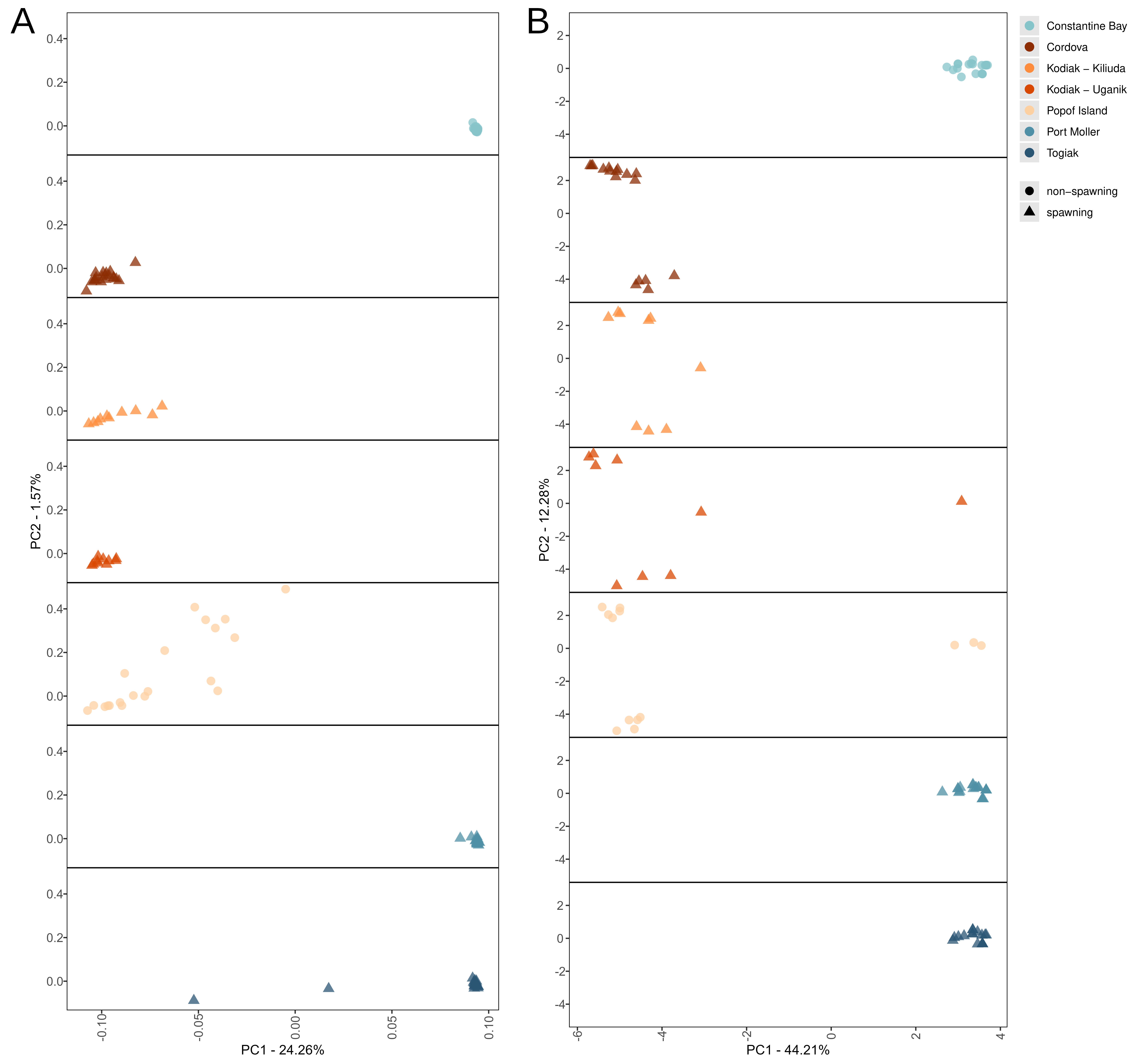

### Fig. S2

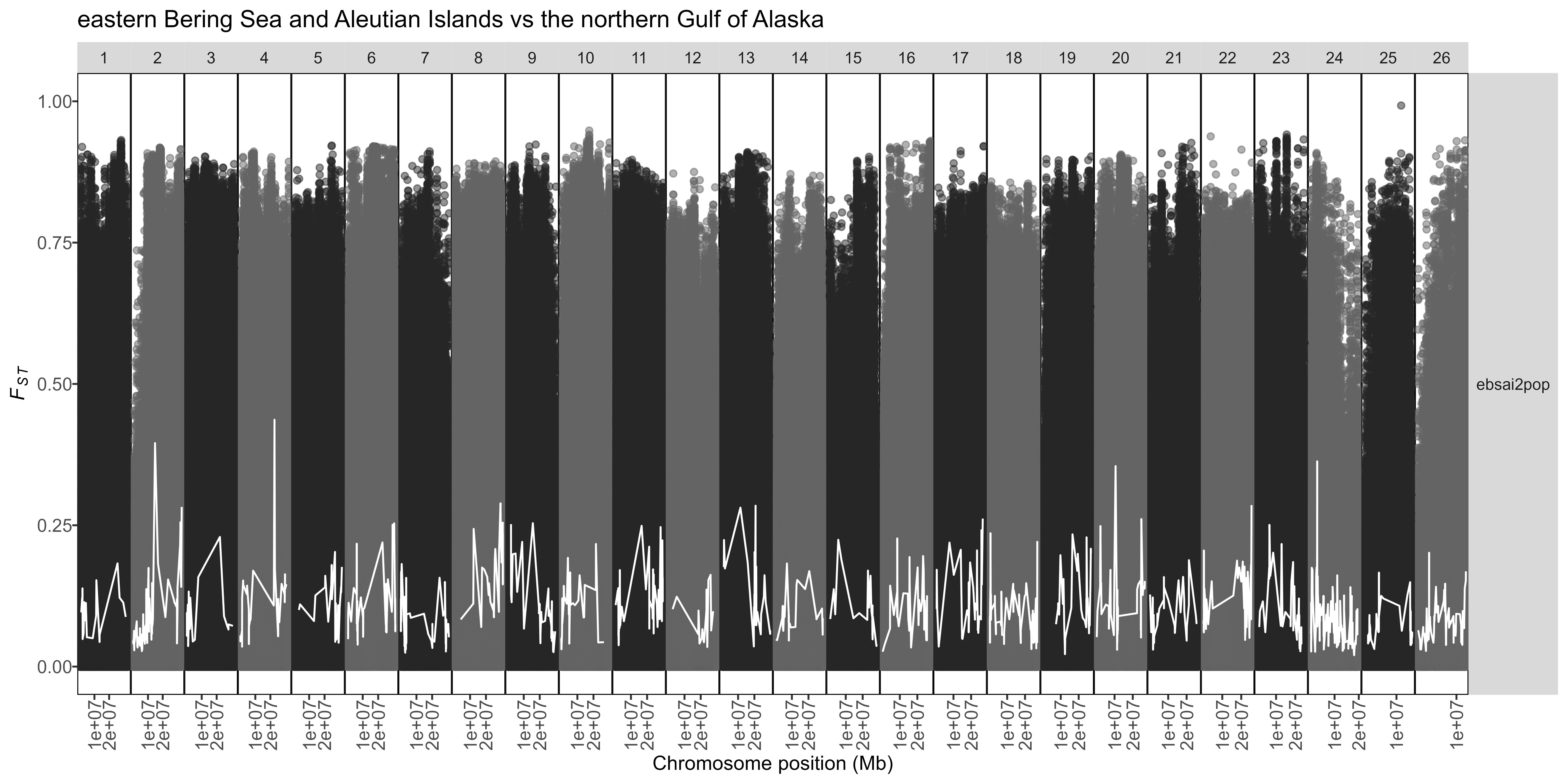

### Fig. S3

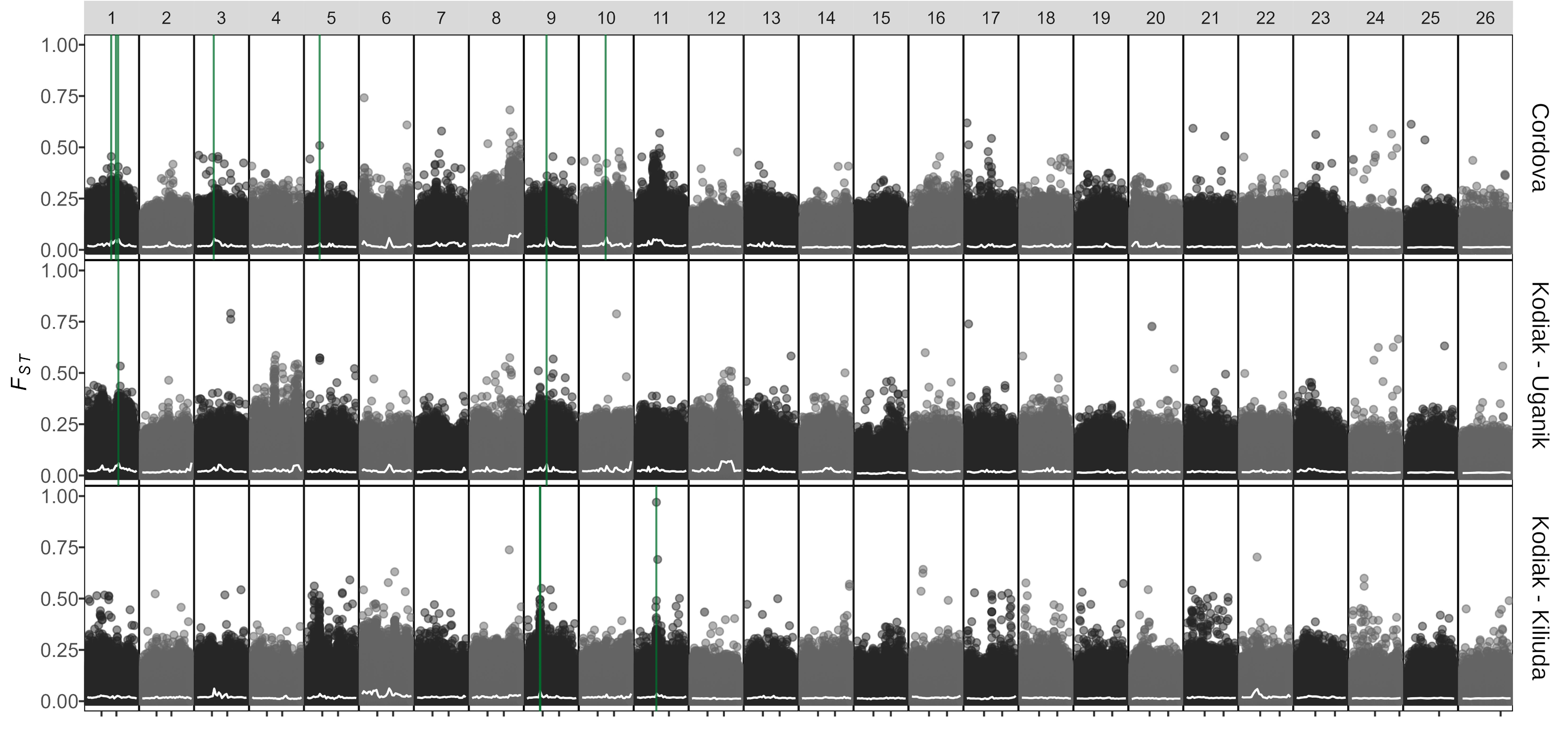

### Fig. S4

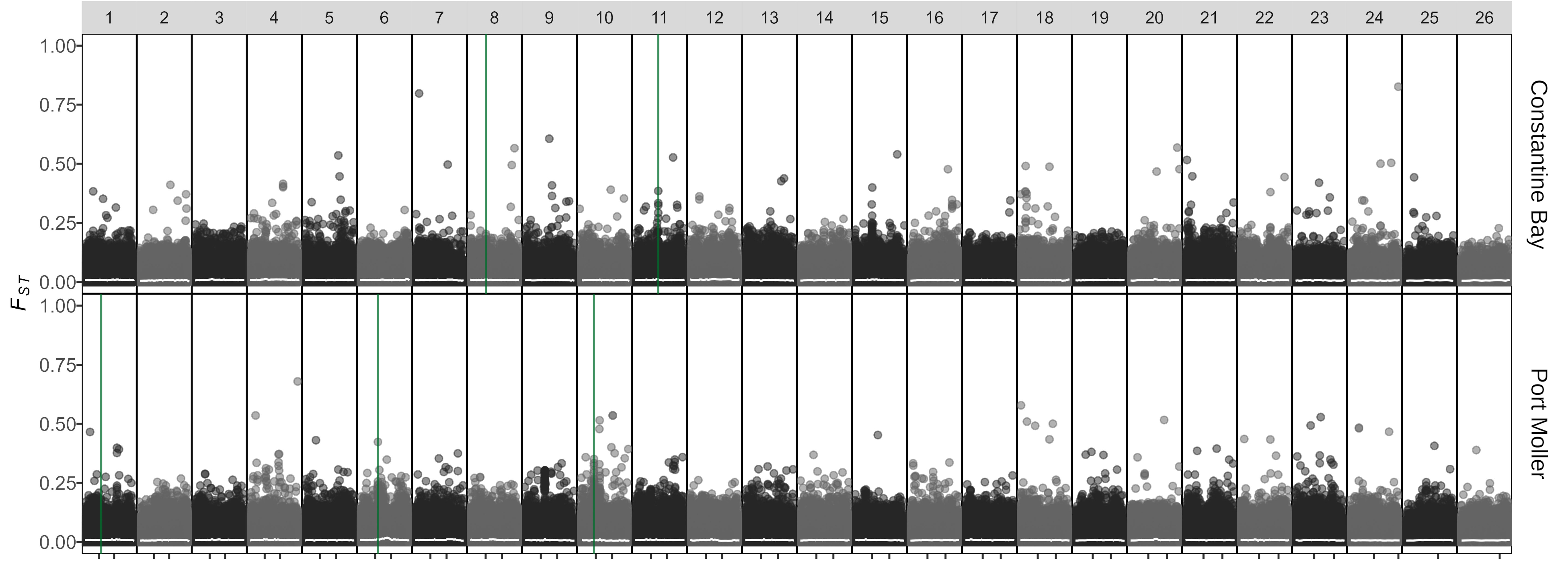

### Fig. S5

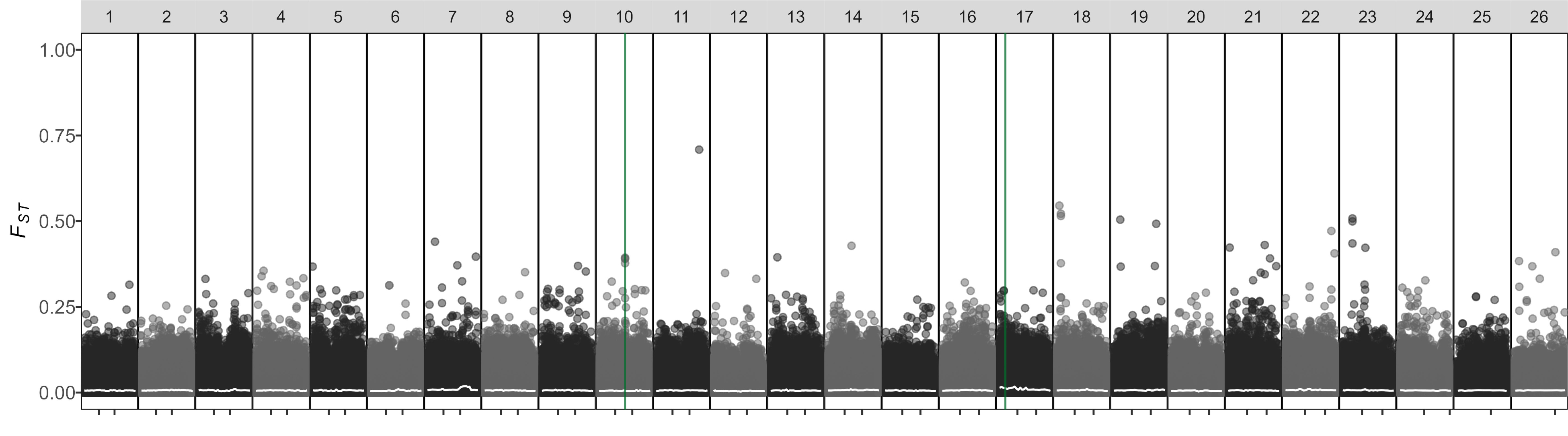
